# Structural dynamics of adenine nucleotide potentiation of the human type 2 IP_3_ receptor

**DOI:** 10.64898/2026.01.23.700653

**Authors:** Viktor Belay, Navid Paknejad, Vinay Sapuru, John D. Chodera, Richard K. Hite

## Abstract

Inositol trisphosphate receptors (IP_3_R) are intracellular calcium (Ca^2+^) channels that mediate Ca^2+^ flux from the endoplasmic reticulum (ER) into the cytosol, playing a critical role in Ca^2+^ signaling. IP_3_R activity requires IP_3_ and Ca^2+^ and is potentiated by adenine nucleotides through a poorly understood mechanism. Here, we combined single-particle cryo-electron microscopy and all-atom molecular dynamics simulations to investigate the potentiation of IP_3_Rs by adenine nucleotides. Our structures reveal that ATP and cAMP bind to a conserved site in the juxtamembrane domain, which connects the cytoplasmic IP_3_- and Ca^2+^-binding sites with the transmembrane pore. Molecular dynamics simulations predict that the binding of adenine nucleotides rigidifies the juxtamembrane domain, primarily through the coordination of the adenine base. Consistent with the adenine base being critical for potentiation, mutations that disrupt the interactions with the adenine base perturb Ca^2+^ flux in cells. Taken together, our data suggest that adenine nucleotides potentiate IP_3_R channel activity by rigidifying the juxtamembrane domain to improve coupling between IP_3_ and Ca^2+^ binding and pore opening.

**Significance Statement:** Inositol-1-4-5-trisphosphate receptors (IP_3_Rs) are the main intracellular calcium (Ca^2+^) release channels in non-excitable cells and contribute significantly to intracellular Ca^2+^ release in excitable cells. Regulation of IP_3_Rs by inositol-1-4-5-trisphosphate (IP_3_), adenine nucleotides, and Ca^2+^ is fundamental to both intracellular Ca^2+^ homeostasis and signaling. We show that adenine nucleotides modulate IP_3_R activity by tuning the dynamics of a mechanical fulcrum-like domain, the juxtamembrane domain (JD), which physically couples the IP_3_- and Ca^2+^-binding sites in the large regulatory cytoplasmic domain to the channel pore. Using a combination of structural, computational, and functional studies, we show that adenine nucleotides bind to and restrict the movement of the JD of the human type 2 IP_3_R (hIP_3_R2). We find that that the coordination of adenine nucleotides is primarily driven by hydrophobic interactions with the adenine moieties of the nucleotides and that these interactions are critical for normal hIP_3_R2 function. This work establishes the foundation for further research into the physiological role of adenine nucleotide modulation of IP_3_Rs.

## Introduction

Inositol-1-4-5-trisphosphate receptors (IP_3_Rs) are large tetrameric cation channels expressed in the endoplasmic reticulum (ER) where they mediate the release of luminal Ca^2+^ into the cytoplasm to regulate diverse cellular processes^1,2,3,4^. Numerous signaling pathways converge upon IP_3_Rs such that their dysfunction is associated with significant pathologies^5,6,7–9,10,11,12^. IP_3_R activity is dependent on the second messenger IP_3_, which is generated by plasma membrane phospholipases in response to extracellular stimuli^13,14^. Protein interaction partners^15^, lipids^16^, and adenine-containing molecules^17,18^ have been demonstrated to further tune IP_3_R activity. In this manner, IP_3_Rs integrate both intracellular and extracellular signals to regulate downstream signaling pathways.

Activation of IP_3_R requires binding of IP_3_ and low Ca^2+^ concentrations, whereas high Ca^2+^ concentrations inhibit channel activity^19,20^. Two Ca^2+^ binding sites reside in the large regulatory cytosolic domain: a high-affinity site whose occupancy favors an activated state and a low-affinity binding site whose occupancy stabilizes the inhibited state^21–23^. Regulation of IP_3_R-mediated Ca^2+^ flux by Ca^2+^ thus occurs in a biphasic manner, with activation occurring at low Ca^2+^ concentrations, and inhibition occurring at high Ca^2+^ concentrations^18^. In cells, this biphasic regulation can give rise to an emergent phenomenon whereby intracellular Ca^2+^ concentrations oscillate locally and globally^24,25,26^. Ca^2+^ oscillations encode signals spatiotemporally in their frequency and amplitude that regulate downstream signaling pathways^27,28^.

Adenine nucleotides broadly potentiate IP_3_R activity by increasing channel open probability at a given concentration of Ca^2+^ and IP_3_^17,29^. Modifications to the adenine base have this effect to varying degrees^30^. Furthermore, potentiation does not depend on the hydrolytic activity of the adenine nucleotides^17^ and does not require the presence of magnesium^31^. This suggests that adenine nucleotides can stably bind to IP_3_Rs. Recent structures of type 1 and type 3 IP_3_Rs have identified a conserved adenine nucleotide binding site in the juxtamembrane domain that connects the ligand-binding containing cytosolic domain ligand binding sites with the pore containing transmembrane domain and channel gate^22,23,32^. However, comparison of static structures with and without ATP shed little light on how adenine nucleotides potentiate IP_3_R activity. Here, we combine single-particle cryo-EM analysis, molecular dynamics simulations, and cellular Ca^2+^ imaging to elucidate the role of adenine nucleotides in IP_3_R activity.

## Results

### Architecture of hIP_3_R2

To investigate the mechanisms by which adenine nucleotides influence the gating of human type 2 IP_3_R (hIP_3_R2) and probe potential low-affinity binding sites, we collected cryo-EM images of nanodisc-embedded hIP_3_R2 in the presence of saturating concentrations of ATP (5.7 mM) or cAMP (5.7 mM) (Supplemental Figure 1A-B). To establish baselines for comparison, we also determined structures of hIP_3_R2 in the presence of a saturating concentration of IP_3_ (74 µM) or in a ligand-free condition. Particle images from the four data sets were combined and subjected to iterative hierarchical classification, yielding three four-fold symmetric classes and four asymmetric classes for each data set (Supplemental Figures 1, 2, 3, 4, 5, and 6 and Supplemental Table 1). Tetrameric hIP_3_R2 channels are composed of a transmembrane domain (TMD) and a large cytosolic domain (CD) (Figure 1A). The TMD contains a pore domain, which is sealed closed in the resolved states at a constriction formed by the side chains of Phe2537 and Ile2541, and a voltage-sensor like S1-S4 domain (Supplemental Figure 7A-D). The CD contains two β-trefoil domains (BTF1 and BTF2) that can assemble into octameric structure known as the BTF ring, three armadillo repeat domains (ARM1, ARM2, and ARM3), two sets of structurally interwoven domains which form the central linker domain and the juxtamembrane domain (JD), and a C-terminal four-helix coiled coil (Figure 1A and Supplemental Figure 7A).

**Figure 1:**
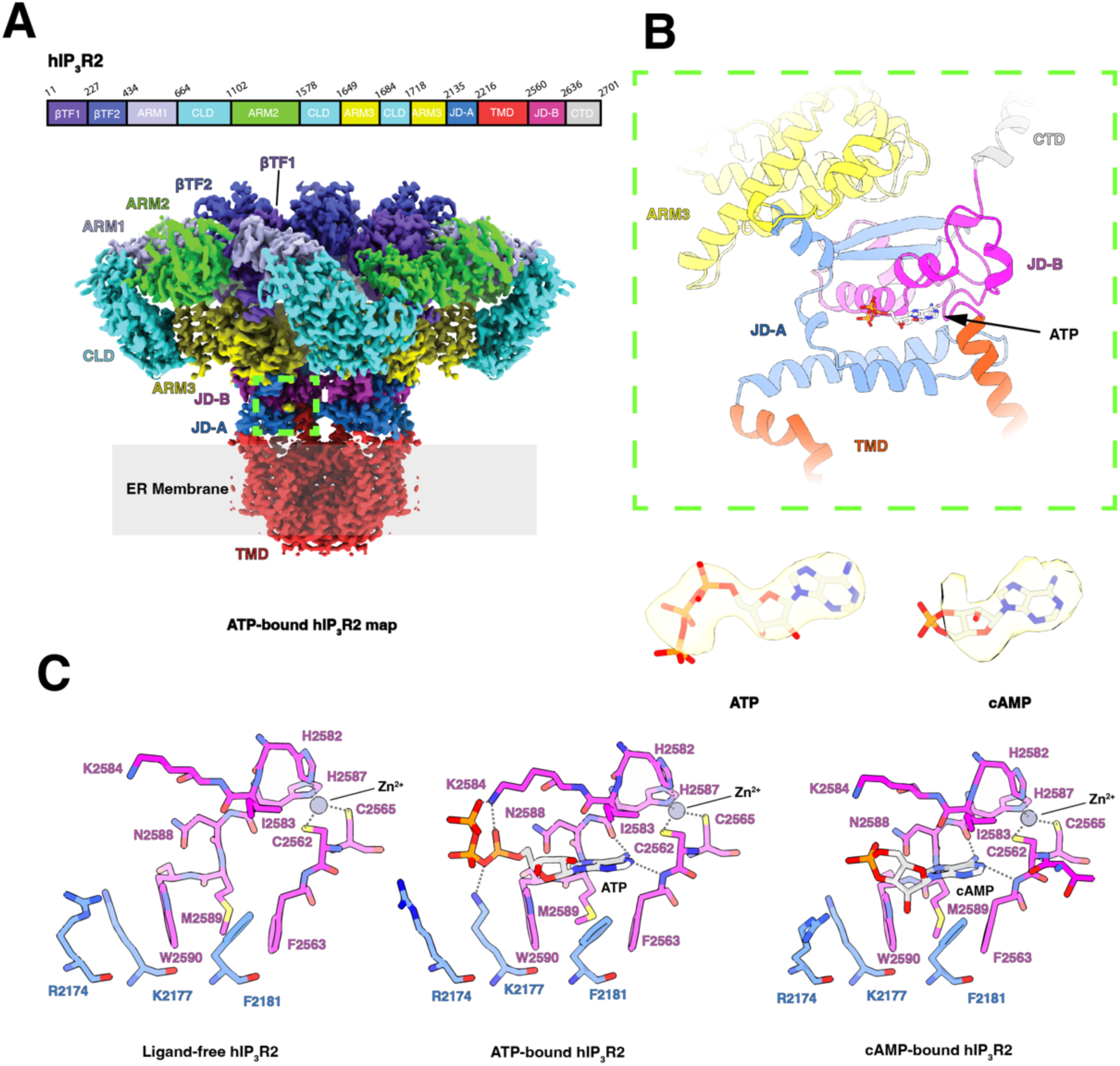
Architecture of hIP_3_R2 and coordination of adenine nucleotides. **A:** Domain organization (top) and composite C4-symmetrized cryo-EM density map of ATP-bound hIP_3_R2 in a resting state, viewed from the side and colored by domain. BTF1 in purple, BTF2 in blue, ARM1 in lavender, ARM2 in green, ARM3 in yellow, CLD in cyan, JD-A in light blue, JD-B in magenta, TMD in red, and CTD in silver. Density for ATP is shown in light yellow. **B:** Structure of the JD and segments of adjacent domains. Cryo-EM densities and fitted atomic models for ATP and cAMP are shown below. Nucleotides are colored by element. **C:** Structure of the hIP_3_R2 adenine nucleotide and Zn^2+^ binding sites in ligand-free, ATP-bound, and cAMP-bound resting states, shown as sticks. JD-A carbons are colored blue, JD-B carbons are colored magenta, and nucleotide carbons are colored gray.

Based on comparisons with previously determined structures of the related type 3 IP_3_R^23^, the seven hIP_3_R2 classes were assigned as an inhibited state, a resting state, a preactivated state, and four resting-to-preactivated transition states. Consistent with its assignment, the inhibited state resembles previous Ca^2+^-bound states in which the BTF ring is disassembled, and the CDs of the four protomers are poorly ordered (Supplemental Figure 5C). Distinguishing the four-fold symmetric resting and preactivated states are the positions of the ARM2 domains, adopting extended conformations in the resting state and retracted conformations in the preactivated state. The resting-to-preactivated transition states represent asymmetric intermediates between the resting and preactivated states in which one two or three of the protomers adopt preactivated-like conformations (Supplemental Figure 6A). The preactivated state was most prevalent among the particles imaged in the presence of IP_3_, which binds at the interface between BTF1 and ARM1 and stabilizes the ARM2 retracted conformation^21–23,32^. A small number of particles imaged in the absence of IP_3_ also adopted the resting-to-preactivated and preactivated states. However, there is only weak density in IP_3_-binding sites (Supplemental Figure 6B), indicating that they are unoccupied or partially occupied by co-purified IP_3_ in these states. These observations suggest that individual hIP_3_R2 protomers can sample the preactivated state spontaneously, and that IP_3_ lowers the energetic barrier between the resting and preactivated states.

Whereas the presence of IP_3_ biased the channel towards the preactivated state, ATP and cAMP did not greatly alter the conformational landscape of hIP_3_R2 compared to the ligand-free condition (Supplemental Figure 5D). Inspection of the resting states derived from the particles imaged in the presence of ATP and cAMP revealed the presence of nucleotides in the adenine nucleotide binding site, whereas the site was empty in the ligand-free resting state and IP_3_-bound preactivated state (Figure 1B, C and Supplemental Figure 5A-B). The adenosine bases of ATP and cAMP adopt similar poses, nestled between the two segments of the JD, which we call JD-A and JD-B, as has been previously observed for ATP binding to type 1 and type 3 IP_3_Rs^22,23,32^. Notably, various adenine nucleotides including ATP have been resolved in the structurally conserved homologous region of Ryanodine Receptors (RyR), a family of distantly related sarcoplasmic reticulum Ca^2+^ channels ^33,34^ (Supplemental Figure 8A-B). In hIP_3_R2, the adenine bases of ATP and cAMP form hydrophobic interactions with Phe2181 of JD-A and Phe2563, Ile2583, and Met2589 of JD-B and hydrogen bonds with Cys2562 and the backbone oxygen of His2587 from the nearby C_2_H_2_ Zn^2+^-finger fold (Figure 1B,C). Densities for the α and β phosphates of ATP are clearly resolved in the density map. The α and β phosphates are directly coordinated by Arg2174 and Lys2177 from JD-A and Lys2584 from JD-B. The γ phosphate of ATP extends away from the binding site and is poorly resolved. Despite being similarly positioned as the α phosphate of ATP, density for the cyclic phosphate of cAMP is not well resolved, indicating that the additional linkage to the ribose limits its ability to interact with hIP_3_R2.

The absence of global conformational changes associated with ATP or cAMP binding indicates that unlike IP_3_ and Ca^2+^, adenine nucleotides do not independently stabilize conformational changes in IP_3_Rs. Previous structural studies investigating IP_3_R activation by IP_3_ and Ca^2+^ revealed that JD-A and JD-B move together as rigid bodies during activation to open the pore and stabilize the activated state^22,23,32^ (Supplemental Figure 9A-B). This movement of the JD directly couples ligand binding in the CD to opening of the pore within the TMD. Combining these observations with our finding that adenine nucleotides are not direct modulators of IP_3_R conformation, we propose that adenine nucleotides potentiate IP_3_R channel activity by stabilizing the interface between JD-A and JD-B to enhance the coupling between the ligand-binding sites in the CD and the pore in the TMD.

### ATP stabilizes the JD

To explore the influence of adenine nucleotides on the conformational landscape of hIP_3_R2, we initiated a series of all-atom molecular dynamics (MD) simulations of the JD of a single protomer of hIP_3_R2 from the ligand-free and ATP-bound resting states (Figure 2A). Since IP_3_R activation occurs on the millisecond timescale^35^ and the JD moves as a rigid body during the activation of the related human type 3 IP_3_R^22,23,32^ (Supplemental Figure 9A-B), we reason that we can faithfully capture the dynamics relevant only to the JD independent of the rest of the protein on the nanosecond to microsecond timescale with a reduced construct comprising JD-A, JD-B, a portion of ARM3, and a portion of the S6 helix. To identify residues in whose position best discriminates between the ligand-free and ATP-bound structures, we computed the absolute difference in root mean square fluctuation (|ΔRMSF|) of the backbone atoms from ligand-free and ATP-bound trajectories in the JD. Regions with high |ΔRMSF| correspond to residues undergoing large ligand-dependent positional change. This analysis revealed residues within the helices that flank ATP, α103 on JD-A and α108 on JD-B, have the highest |ΔRMSF| among structured elements in the JD (Supplemental Figure 10A-B).

**Figure 2:**
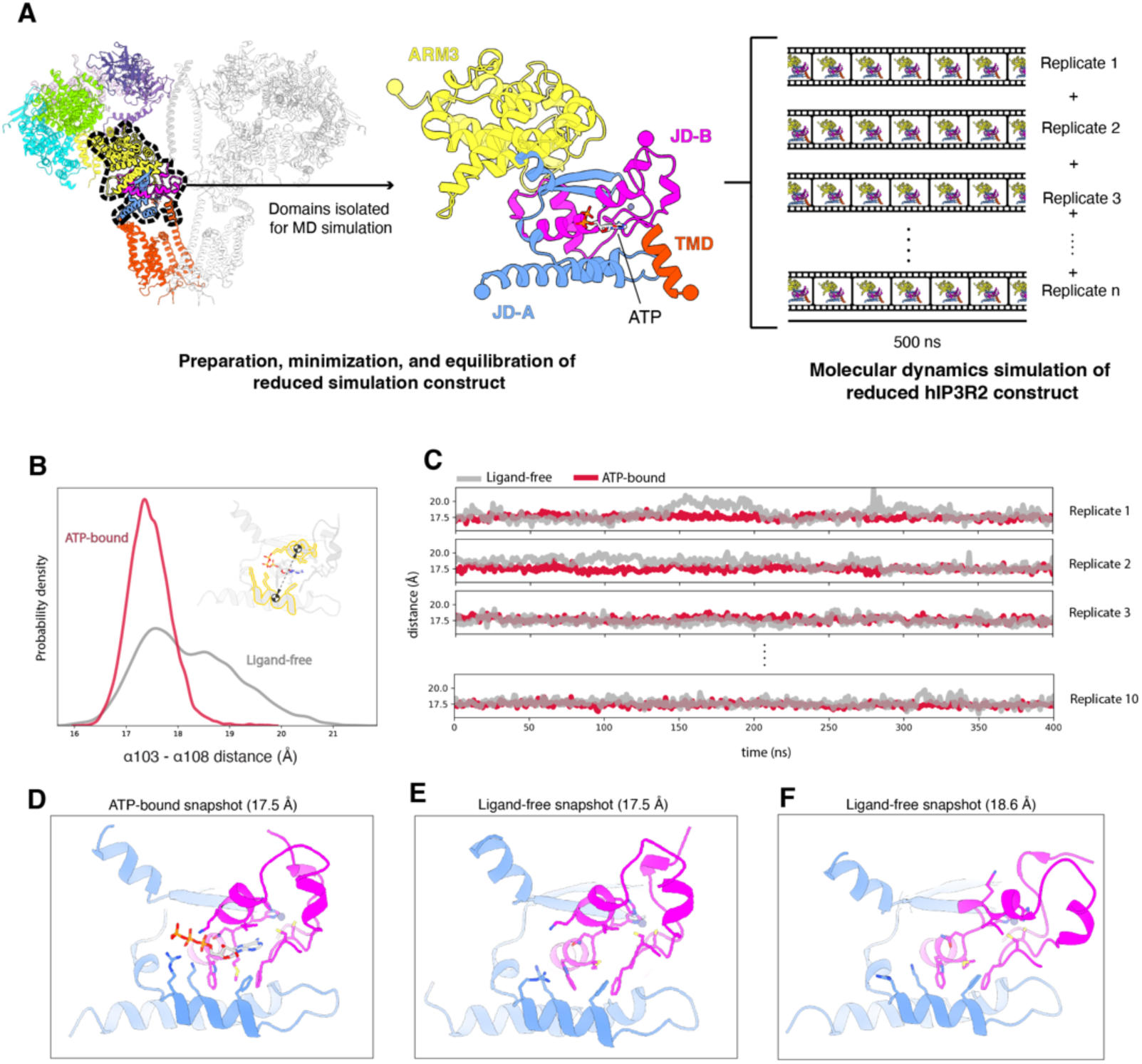
ATP stabilizes the JD-A/JD-B interface in MD simulations. **A:** Schematic depicting the construction and simulation of the hIP_3_R2 MD construct used throughout this study. JD-A, JD-B, a portion of ARM3, and a portion of the TMD of a single hIP_3_R2 protomer are extracted from the cryo-EM structure of resting hIP_3_R2, placed in a water box with neutralizing NaCl, energy minimized, and equilibrated. Ten production simulations of the minimized and equilibrated construct proceeded for 400 ns for a total aggregate simulation time of 4 µs for each condition. **B:** Kernel density estimation (KDE) of the distance between the centers of mass (COM) of the backbone atoms of α103 and α108 from either ligand-free trajectories (gray) or ATP-bound trajectories (red). A black dashed line and COM markers indicate the measured distances in the inset. The residues included in the measurement are highlighted in gold on the atomic model of the JD. **C:** Representative traces of the α103-α108 backbone COM distance measurement from simulation replicates of ligand-free trajectories (gray) and ATP-bound trajectories (red). **D-F:** Structural snapshots of JD-A and JD-B with either ATP bound or ligand-free from MD simulations. **D:** Snapshot with ATP bound where the α103-α108 COM distance is 17.5 Å. **E:** Snapshot with no ligand bound where the α103-α108 COM distance is 17.5 Å. **F:** Snapshot with no ligand bound where the α103-α108 COM distance is 18.6 Å

Based on the results of the |ΔRMSF| analysis, we next evaluated the effect of ATP on the conformation of the nucleotide-binding site during the simulations by measuring the distance between the center of mass of the backbone atoms of α108 and the center of mass of the of the backbone atoms of the adjacent helix α103 in all frames of the ATP-bound and ligand-free trajectories. We find that the distribution of distances in the ATP-bound trajectories is unimodal with a mean distance of 17.48 ± 0.01 Å (Figure 2B, Supplemental Table 2). The distribution is tight around its mean, with a standard deviation (SD) of 0.4 Å. In contrast, the distribution of the ligand-free distances shows a multimodal distribution with an overall mean distance of 18.13 ± 0.06 Å and SD of 1.0 Å. Fitting two gaussians to the ligand-free distance distribution shows the first peak having a mean distance of 17.6 Å and the second having a mean of 19.0 Å (Supplemental Figure 11A). Although the first peak partially overlaps with the ATP-bound distribution, the second peak is distinct, indicating that the second peak corresponds to structural configurations not sampled by the adenine nucleotide bound systems (Figure 2B). Thus, ATP restricts the diversity of configurations that can be sampled by the JD.

Upon inspection of representative snapshots of the JD around the mean of the ATP-bound distance distribution, we find that the JD adopts a conformation closely resembling the cryo-EM structure from which the trajectories initiated (Figure 2C). Consistent with the bimodal distribution of distances, snapshots taken from the ligand-free trajectories reveal multiple distinct states. Representative snapshots from the first peak resemble the initiating cryo-EM structure whereas the nucleotide-binding site is distorted in snapshots taken from the second peak (Figure 2E-F). For example, in a ligand-free snapshot in which the distance between helices α103 and α108 is 18.6 Å, the side chain of Arg2174 is oriented away from the binding site and helix α108 is partially unwound, resulting in a more than 10.0 Å movement of Lys2584. Together, the outward movements of Arg2174 and Lys2584 open the binding pocket and alter its electrostatic potential. Indeed, there is a significant correlation between the distance between Arg2174 and Lys2584 and the distance between helices α103 and α108 in the ligand-free trajectories (Supplemental Figure 11B), indicating that position of helices α103 and α108 inform on the conformation of the nucleotide-binding site as well as on the JD as a whole.

### The adenine base is critical for stabilizing the JD

To further interrogate the interactions between the JD and nucleotides, we replaced the ATP in the ATP-bound structure with ADP, AMP, adenosine or guanosine and performed additional MD simulations. We also initiated MD simulations from the cryo-EM structure of the cAMP-bound resting state. To quantify the effects of nucleotides on the conformation of the JD, we measured the distance between the centers of mass of the backbone atoms of helices α103 and α108. The distance distributions for the ADP, AMP, cAMP, and adenosine trajectories were unimodal with mean distances nearly identical to those observed in the ATP-bound trajectories whereas the distribution for the guanosine trajectories was bimodal (Figure 3A). The overall conformation of the JD in representative snapshots of trajectories performed with ADP, AMP, cAMP, and adenosine were indistinguishable from those performed with ATP (Supplemental Figure 11C). Moreover, the adenosine base remained stably bound in the nucleotide-binding site and adopted a similar pose as did the adenosine base of ATP in the ATP trajectories. In contrast, the guanosine base was rapidly ejected in all of the trajectories, consistent with previous binding experiments that demonstrated that guanine nucleotides have minimal impact on IP_3_R (Figure 3B,E-F)^30,36^.

**Figure 3:**
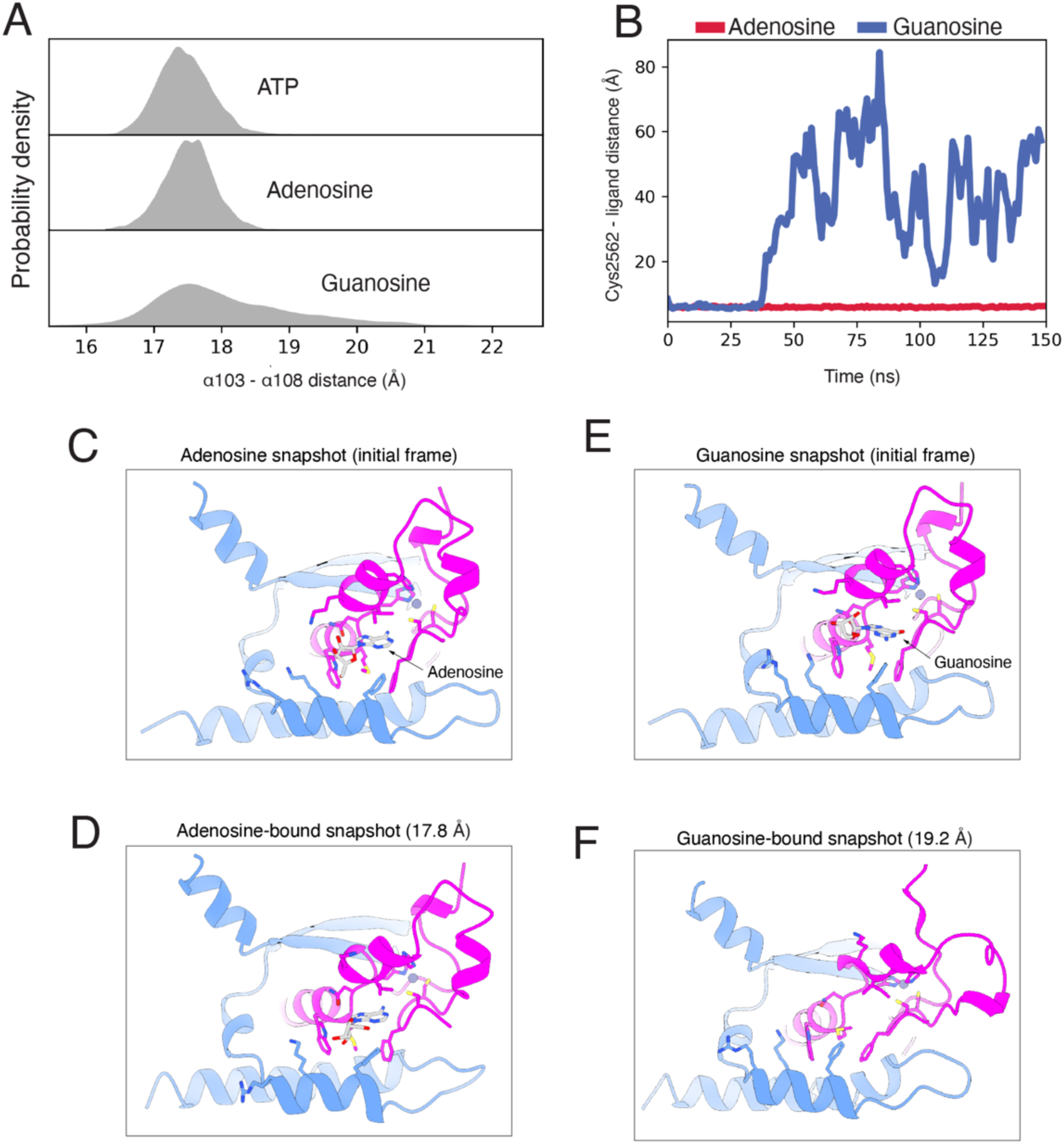
Adenosine is minimally required to stabilize the JD-A/JD-B interface. **A:** KDEs of the α103-α108 COM distance from ATP-bound, adenosine-bound, or guanosine-bound trajectories. ATP-bound α103/α108 COM distance distribution is the same as presented in figure 2. **B:** Representative time series of the distance between either adenosine (red line) or guanosine (blue line) and cys2562. **C:** Structural snapshot of JD-A and JD-B with adenosine bound from the initial simulation frame. **D:** Structural snapshot of JD-A and JD-B with adenosine bound from simulations of a frame where the α103-α108 COM distance is 17.8 Å. **E:** Structural snapshot of JD-A and JD-B with guanosine bound from the initial simulation frame. **F:** Structural snapshot of JD-A and JD-B from a simulation that started with guanosine bound after guanosine was ejected from the binding site. The α103-α108 COM distance in this frame is 19.2 Å.

Although the backbone distance distributions for all adenine-containing trajectories were similar to those bound with ATP, differences emerged in side-chain conformation and occupancy. Arg2174, Lys2177, and Lys2584 displayed increased flexibility in simulations with adenine nucleotides other than ATP, showing a stronger propensity to sample solvent-facing conformations in simulations with shorter phosphate tails (Supplemental Figure 11C-D). Together, these observations suggest that the adenine base, through its interactions with the two segments of the JD, rigidifies the JD, whereas the phosphate groups play accessory roles by tuning local side-chain dynamics.

### Perturbing the adenine nucleotide binding site disrupts cellular Ca^2+^ oscillations

To assess the role of the interactions between JD and adenosine nucleotides, we perturbed the adenine nucleotide binding site in hIP_3_R2 and quantified the effects on Ca^2+^ release dynamics in intact cells. We generated *hITPR2-eGFP* fusions in which we mutated Arg2174, Lys2177, or Lys2584, which interact with the phosphate groups, or Met2589, which interacts with the adenine base. We first assessed the effect of the mutations on hIP_3_R2 folding by transducing these constructs into IP_3_R-null HEK293T cells, solubilizing proteins with detergent and then monitoring their migration using fluorescence detection size exclusion chromatography. The wild-type and mutant constructs all eluted at ∼11.5 mL with similar peak heights, indicating that the mutations do not alter tetrameric assembly and have minimal effect on protein expression (Supplemental Figure 12A-B).

We next incubated IP_3_R-null HEK293T cells transduced with either wild-type or mutant *hITPR2-eGFP* fusion constructs with the membrane permeable Ca^2+^-sensitive dye Calbryte 590 and stimulated IP_3_ production by perfusing cells with a saturating concentration (100 uM) of the muscarinic acetylcholine receptor agonist carbachol (Figure 4A). We monitored cells for hIP_3_R2-dependent intracellular Ca^2+^ release using live-cell fluorescence imaging. Carbachol stimulated oscillatory Ca^2+^ responses in IP_3_R-null cells transduced with wild-type hIP_3_R2, but not in IP_3_R-null cells (Figure 4B,D). Oscillatory responses occurred in 76 percent of the cells expressing hIP_3_R2 and occurred with a mean of 2.3 peaks per cell for cells that had an oscillatory response. Cells expressing mutants predicted to disrupt interactions with the phosphate groups behaved comparably to cells expressing wild-type hIP_3_R2, with oscillatory responses observed in 69 percent of the cells expressing the triple alanine mutation (R2174A/K2177A/K2584A) and 73 percent of the cells expressing the triple glutamate mutation (R2174E/K2177E/K2584E). Conversely, only 27 percent of the cells expressing M2589R and 24 percent of the cells expressing M2589W yielded oscillatory responses (Figure 4D). These data indicate that the coordination of the adenine base within the JD is critical for activation of IP_3_Rs whereas interactions with the phosphate groups are not required for oscillatory response in cells under saturating carbachol stimulation. Moreover, these results are consistent with our MD simulations which predict that interactions between adenine nucleotides and their stabilization effect on the JD are primarily driven by the adenine moiety.

**Figure 4:**
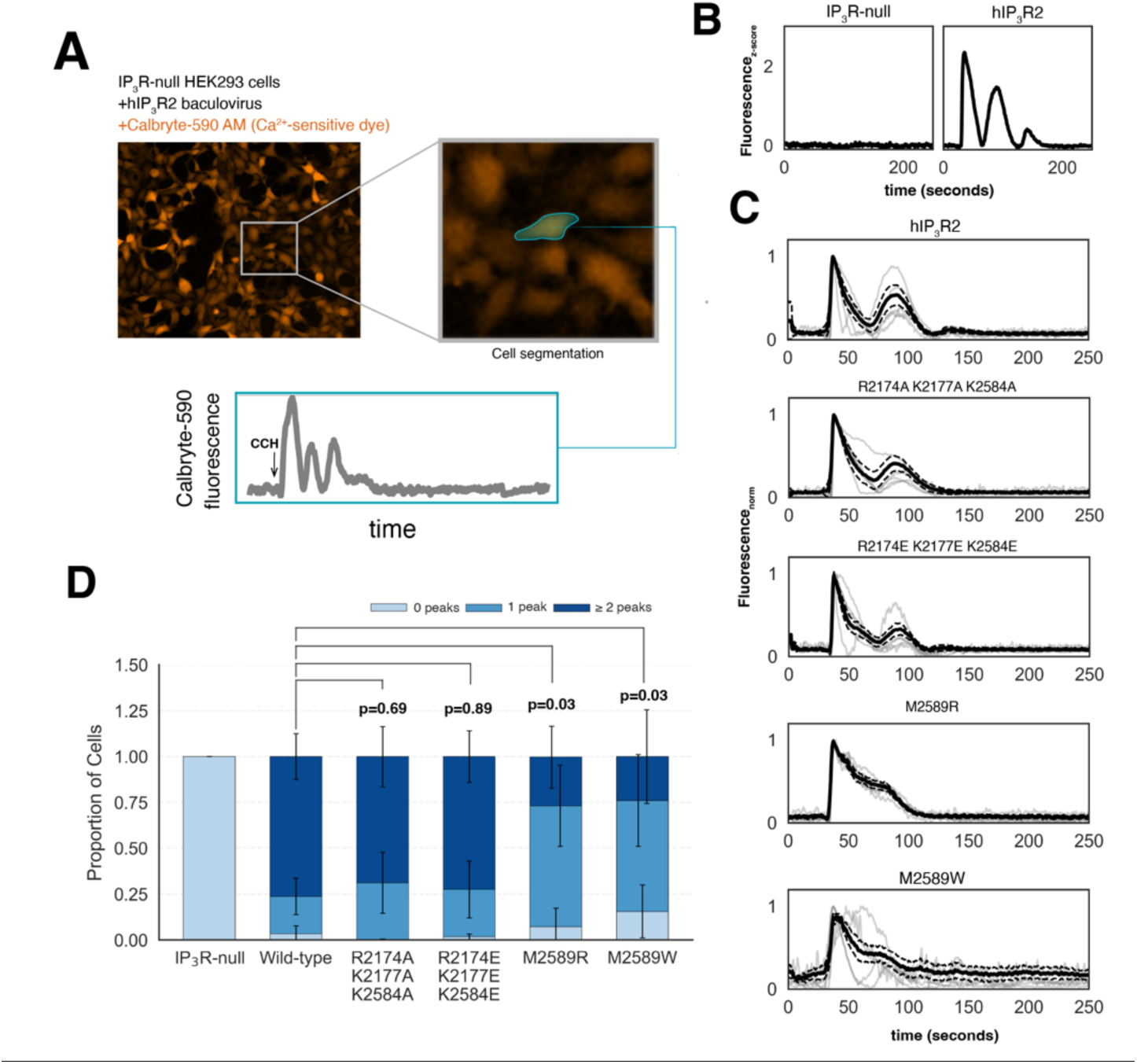
Disrupting interactions with the adenine base of ATP diminishes Ca^2+^ oscillations in cells. **A:** Schematic of cellular Ca^2+^ imaging assay. (Top) Representative fluorescence image of IP_3_R-null HEK293T cells expressing hIP_3_R2 with inlet showing individual segmented cells. (Bottom) Representative Ca^2+^ fluorescence trace from a single segmented cell over time. Cells were stimulated with carbachol (CCH) at the time highlighted by the arrow. **B:** Representative z-score normalized time-course traces of Ca^2+^ fluorescence recorded from an IP_3_R-null cell (left) or a cell expressing wild-type hIP_3_R2 (right). **C:** Representative time course traces of Ca^2+^ fluorescence recorded from IP_3_R-null cells expressing either wild-type or mutant hIP_3_R2. Solid black line represents mean of twenty randomly selected cells from one representative plate that yielded at least one peak. Dashed black lines represent the 95% confidence interval around the mean trace. Translucent gray lines are six of the twenty traces, selected randomly. Traces are aligned by the first peak and normalized between 0 and 1. **D:** Distribution of Ca^2+^ responses in non-transduced IP_3_R-null cells or IP_3_R-null cells transduced with either wild-type or mutant hIP_3_R2 in response to carbachol stimulation. Stacked bars represent the mean proportion of cells exhibiting zero, one, or two or more Ca^2+^ peaks across four plates for each construct. Error bars represent the standard deviation in the mean proportion for each response category. P-values indicate statistical significance between proportions of wild-type and mutant cells with two or more peaks (calculated by Mann-Whitney U-test).

To visualize how these mutants shape JD stability, we next introduced the same mutations *in-silico,* performed MD simulations of the isolated JD ATP-bound construct, and measured the distance between the centers of mass of backbone atoms of helices α103 and α108 for each mutant construct. The distance distribution for the triple alanine mutation was unimodal and only slightly shifted rightward compared to the wild-type trajectories, with the mean distance between the backbone atoms of helices α103 and α108 being 17.68 ± 0.01 Å. The distribution of the triple glutamate mutation was shifted the distribution rightward to a mean of 18.13 ± 0.03 Å and displayed a slight bimodality (Figure 5A, Supplemental Table 2). Despite the appearance of a secondary mode in the triple glutamate mutation, the distance distribution of the triple glutamate construct displayed a narrower spread than the ligand-free construct, indicating that the JD samples a restricted conformational landscape even with the triple glutamate mutation. The distance distributions of the M2589W and M2589R mutants were shifted rightwards to mean distances of 18.28 ± 0.13 Å and 19.36 ± 0.08 Å, respectively (Figure 5A, Supplemental Table 2). The shifts were accompanied by the appearance of additional modes and a broadening of the distance distribution, indicating the presence of alternative conformations. Indeed, representative snapshots of alternative modes from trajectories of the M2589W and M2589R mutants reveal distortions in the nucleotide-binding site as well as more global rearrangements of the JD (Figure 5F-I). Thus, the computational and functional analyses are in accordance, both highlighting the critical role of adenine bases in the activation of IP_3_Rs.

**Figure 5:**
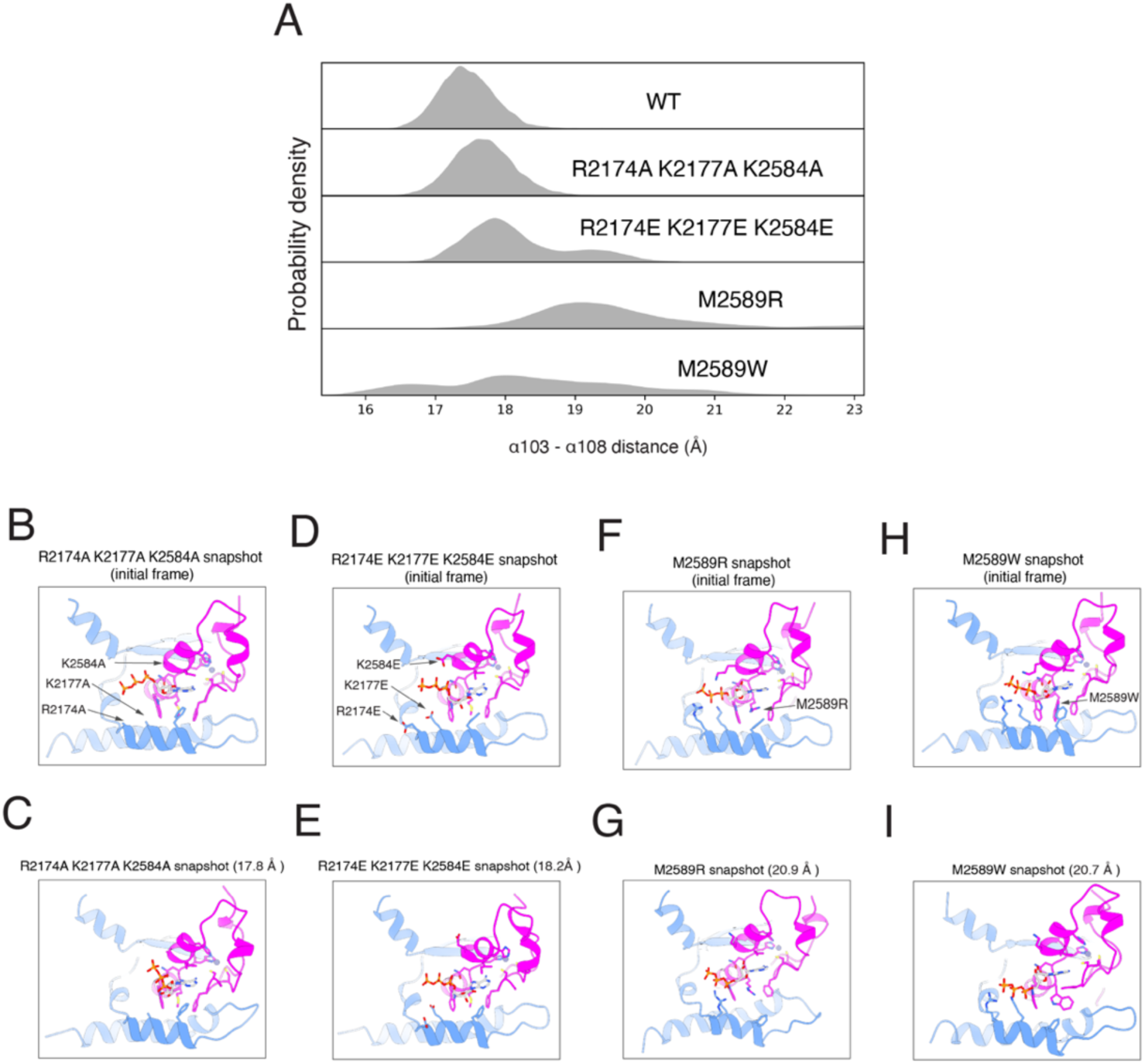
Mutations to the adenine binding region disrupt stability in the JD-A/JD-B interface in the ATP-bound JD in MD simulations. **A:** KDEs of the α103-α108 COM distance in the ATP-bound state for wild-type hIP_3_P2, R2174A K2177A K2584A, R2174E K2177E K2584E, M2589R, or M2589W. The ATP-bound α103/α108 COM distance is the same as presented in figure 2. **B-C:** Initial (B; α103-α108 COM distance = 16.9 Å) and representative (C; α103-α108 COM distance = 17.8 Å) structural snapshots of JD-A and JD-B from simulations of the triple mutation R2174A, K2177A, and K2584A bound with ATP. **B:** Snapshot at the initial frame **C:** Snapshot where the α103/α108 COM distance is 17.6 Å. **D-E:** Initial (B; α103-α108 COM distance = 17.9 Å) and representative (C; α103-α108 COM distance = 18.2 Å) structural snapshots of JD-A and JD-B from simulations of the triple mutation R2174E, K2177E, and K2584E bound with ATP. **D:** Snapshot at the initial frame **E:** Snapshot where the α103/α108 COM distance is 18.2 Å. **F-G:** Initial (B; α103-α108 COM distance = 18.4 Å) and representative (C; α103-α108 COM distance = 20.9 Å) structural snapshots of JD-A and JD-B from simulations of the mutation M2589R bound with ATP. **F:** Snapshot at the initial frame **G:** Snapshot where the α103/α108 COM distance is 20.9 Å. **H-I:** Initial (B; α103-α108 COM distance = 17.9 Å) and representative (C; α103-α108 COM distance = 20.7 Å) structural snapshots of JD-A and JD-B from simulations of the mutation M2589W bound with ATP. **H:** Snapshot at the initial frame **I:** Snapshot where the α103/α108 COM distance is 20.7 Å.

## Discussion

Here, we investigated the role of adenine nucleotides in the activation of IP_3_Rs, finding that the adenine nucleotides act as molecular glues that stabilize the interface between the two segments of the JD. Located at the interface between the IP_3_- and Ca^2+^-binding sites in the CD and the pore in the TMD, the JD participates in the long-range allosteric activation of IP_3_Rs. Upon IP_3_ and Ca^2+^ binding, conformational changes in the CD are transmitted through the JD to open the pore. Pore opening involves both connections of the JD to the TMD, with JD-A reorienting the S1-S4 domain and JD-B pulling the pore-lining S6 helices apart^22,23,32^. By rigidifying the two segments of the JD, adenine nucleotides enable the JD to faithfully transduce the conformational changes in the CD to both domains of the TMD and open the pore.

The rigidification of the JD is primarily driven by the adenine base, which is sandwiched between the two segments of the JD through hydrophobic interactions. In the absence of an adenine nucleotide, the simulations reveal that the interface between JD-A and JD-B can expand, which would weaken the association between the two segments. The phosphate groups, which interact with nearby positively charged residues, have lesser effects. Simulations with nucleotides with shorter phosphate groups displayed increased side chain dynamics rather than larger-scale alterations in the JD. Consistent with these results, mutations that disrupt phosphate tail coordination have no detectable effect on hIP_3_R2 activity in cells, whereas mutations that disrupt adenine coordination significantly diminished activity. Together, these findings suggest that adenine nucleotide binding rigidifies the JD and that potentiation occurs primarily via adenine-mediated stabilization.

More generally, adenine nucleotides may act to couple diverse regulatory interactions in the CD with the channel gate. Recent work has shown that ATP can also influence channel inhibition ^37^ possibly also by facilitating the long-range coupling of inhibitory signals from the CD to the gate. Overall, our model offers a unifying structural framework for how adenine nucleotides may tune IP3R activity under different regulatory regimes.

Ryanodine receptors (RyR) are a family of sarcoplasmic reticulum Ca^2+^ release channels that are distantly related to IP_3_Rs. RyRs possess a domain centered on the Thumb and Forefingers (TaF) domain and the C-Terminal domain (CTD) that is structurally homologous to the JD of IP_3_Rs (Supplemental Figure 8A-B)^38^. Like the IP_3_R JD, the TaF domain and CTD together couple the large RyR cytoplasmic domain with the transmembrane domain and contain a conserved adenine nucleotide binding pocket (Supplemental Figure8)^33,34,39^. Although adenine nucleotides have different effects on RyRs compared to IP_3_Rs, potentially including direct activation of RyRs, the potency of adenine nucleotides towards RyRs is also graded depending on phosphate tail length (ATP > ADP > AMP) ^40,41^. Recent structural work suggests that ATP strengthens the interaction at the TaF domain/CTD interface, reinforcing force transmissions between the large cytoplasmic domain and the channel gate^42^. This parallels our finding that adenine nucleotides rigidify the IP_3_R JD to strengthen long-range allosteric signaling. Together, these structural and functional similarities suggest a shared mechanism by which adenine nucleotides stabilize a conserved domain which facilitates transmission of long-range mechanical signals.

The precise physiological role of adenine nucleotide binding to IP_3_Rs remains unclear. Intracellular ATP exists largely as Mg-ATP (ATP^2-^) at a concentration of ∼1-5 mM^43^ and therefore would likely be constitutively bound to IP_3_Rs based on previously determined binding affinities^36^. However, Mg^2+^ is not required for IP_3_R modulation by ATP and sequence differences between the three IP_3_R isoforms may alter the effect of Mg^2+^ on ATP coordination ^17,31^. Whereas the phosphate tail in type 1 and type 2 IP_3_Rs is coordinated by lysine and arginine residues, a glutamate residue is present in type 3 that may allow coordination of a Mg^2+^ ^32^. It is thus possible that free ATP (ATP^4-^), which exists at a concentration of ∼100 uM in cells^44,45^, may contribute to the modulation of IP_3_R activity in cells. Since IP_3_Rs natively form heteromeric assemblies^46,47^, varied tetrameric isoform composition could tune IP_3_R sensitivity to ATP and other adenine nucleotides in the cell. Future work is necessary to establish the identity of the adenine nucleotides responsible for IP_3_R modulation and the physiological consequence of this regulation.

Altogether, our study contributes to the broader understanding of ligand regulation and mechanisms of allosteric action within IP_3_R. By combining structural information from cryo-EM, protein dynamics obtained from MD simulations, and cell-based functional measurements, we have been able to generate and test hypotheses about the structural determinants of adenine nucleotide potentiation of IP_3_R. We have demonstrated how integrating these approaches can provide deeper insights into how ligands modulate activity in large, highly allosteric proteins such as IP_3_R.

## Methods

### Suspension cell culture

For protein expression, human embryonic kidney (HEK293S) cells lacking N-acetylglucosaminyltransferase I (GnTI^-^) were acquired from ATCC. HEK293S GnTI^-^cells were maintained in suspension in FreeStyle 293 Expression Medium (Fisher Scientific) supplemented with 5% FBS with constant agitation at 225 RPM at 37°C with 8% CO2.

For baculovirus production, *Spodoptera frugiperda* (sf9) cells were acquired from Expression Systems. sf9 cells were maintained in suspension in serum-free insect cell media (Expression Systems) supplemented with 5% FBS and 1% penicillin/streptomycin with constant agitation at 225 RPM at 27°C with 0% CO2.

### Molecular biology

The *hIP_3_R2* gene was synthesized into a plasmid containing XhoI and EcoRI sites for subcloning. The *hIP_3_R2* gene was linked to a C-terminal *eGFP* via a human rhinovirus 3 C protease cut-site^48^. Point mutations were made to *hIP_3_R2* via HiFi DNA assembly (NEB). Oligonucleotides were designed using the NEB primer design tool (https://nebuilder.neb.com/) to generate two overlapping plasmid fragments with one of the fragments containing nucleotide substitutions intended to induce site-directed mutagenesis. Oligonucleotides used for mutagenesis are listed in Supplemental Table 3. All constructs were verified by whole plasmid sequencing performed by Plasmidsaurus using Oxford Nanopore Technology with custom analysis and annotation.

Constructs were used to transform DH10Bac *Escherichia coli* cells to generate bacmids. Bacmids were transfected into sf9 cells to generate P1 recombinant baculovirus. 100 µg of purified bacmid was incubated with 400 µg of 25,000 MW polyethyleneimine (Polysciences) in Milli-Q water at a final volume of 1400 µL. This solution was incubated at 50°C for 30 minutes to sterilize. This solution was then added to 50 mL of sf9 cells at 10^6^ cells/mL grown in suspension at 27°C and cultured in serum-free insect cell media (Expression Systems) supplemented with 5% fetal bovine serum (FBS) and 1% penicillin/streptomycin. Viral titer was amplified to P3 and was separated from cell debris by centrifugation at 3800 RPM for 10 minutes.

### hIP_3_R2 expression and purification

200 mL of P3 baculovirus containing wild-type *hIP_3_R2* was used to transduce 4 L of HEK293S GnTI^-^ cells. Valproic acid at a final concentration of 3.75 mM was added at the time of transduction. Transduced cells were harvested 72 h following transduction by centrifugation at 4000 RPM for 20 minutes, washed with phosphate buffered saline (PBS) without Ca^2+^ or Mg^2+^, and flash frozen.

Cell pellets were lysed in a solubilization buffer with 27.5 mM CHAPS, 300 mM NaCl, 20 mM HEPES pH 7.5, 1 mM phenylmethylsulfonyl fluoride (PMSF), 2.5 µg/mL aprotinin, 2.5 µg/mL leupeptin, 1 µg/mL pepstatin A, 0.5 mM 4-benzenesulfonyl fluoride hydrochloride (AEBSF), 1 mM benzamidine, and a few flakes of deoxyribonuclease I for 2 h at 4 C. The cell lysate was centrifuged for 50 minutes at 29,000×g and the supernatant was then incubated with GFP-affinity beads for 4 h at 4 C with constant inversion. The beads were collected on a column and washed with 100 mL of a buffer containing 4 mM CHAPS, 450 mM NaCl, 50 mM Tris-HCl pH 8.0, 10 mM EGTA, 10 mM HEDTA, 2 mM dithiothreitol, 0.075 mg/mL lipids 4:1:1:1=POPC:POPE:POPS:brain Extract. The beads were then harvested by centrifugation at 1000 RPM for 5 min and excess buffer was removed by pipetting. 200 uL of lipids (10 mg/mL of 4:1:1:1=POPC:POPE:POPS:brain Extract solubilized in 35 mM CHAPS, 450 NaCl, 50 mM Tris-HCl pH 8.0) and 400 uL of 5 mg/mL MSP1E3D1 were added to the beads and this solution was incubated with constant inversion for 30 min. BioBeads were added to double the volume of this slurry, and the slurry was incubated with constant inversion for 8 h. The GFP-affinity beads and bioBeads mixture was collected on a column and washed with 100 mL of 450 mM NaCl, 50 mM Tris-HCl pH 8.0, 10 mM EGTA, 10 mM HEDTA, and 2 mM DTT. The bead mixture was then incubated with 3C protease diluted in wash buffer on the column for 4 h to cleave hIP_3_R2 from the affinity resin. The elution was concentrated to 250 µL and further purified with size exclusion chromatography (SEC) on a Superose 6 Increase column (GE Life Sciences) in a buffer containing 100 mL of 450 mM NaCl, 50 mM Tris-HCl pH 8.0, 10 mM EGTA, 10 mM HEDTA, and 2 mM DTT. Peak SEC fractions were collected, and the protein was concentrated to 15 mg/mL.

### Electron microscopy sample preparation

For the apo hIP_3_R2 condition, 3 uL of 15 mg/mL purified protein was incubated with 2.3 mM fluorinated fos-choline 8 (FIFC8) for 1.5 h on ice and applied to glow-discharged gold 400-mesh Quantifoil R1.2/1.3 holey carbon grids (Quantifoil). Grids were plunged into liquid-nitrogen-cooled liquid ethane using a Vitrobot Mark IV (Thermo Fisher). For liganded conditions, 3 uL of 15 mg/mL purified protein was incubated with 2.9 mM FIFC8 and either 74 µM IP_3_, 5.7 mM ATP, or 5.7 mM cAMP for 1.5 h on ice. Samples were then applied to glow-discharged gold 400-mesh Quantifoil R1.2/1.3 holey carbon grids and the grids were plunged into liquid-nitrogen-cooled liquid ethane using a Vitrobot Mark IV.

### Electron microscopy data acquisition, analysis, and model building

Cryo-EM images were collected at 0.826 Å/pixel on a Titan Krios with a total dose of 66 e^-^/Å^2^ at the Pacific Northwest Cryo-EM Center. One dataset was collected for each ligand condition during the same session, resulting in 5,282 movies for apo, 4,450 movies for +ATP, 7,272 movies for +cAMP, and 7,651 movies for +IP_3_.

Cryo-EM movies from each condition (apo, +IP_3_, +ATP, +cAMP) were combined and processed in CryoSPARC Live version 3.2 for motion correction and contrast transfer function (CTF) estimation. Particles were picked using a blob-picker with a gaussian blob with dimensions between 150 and 250 Å, yielding a stack of 7,961,605 particles. Picked particles were extracted with a box size of 512 × 512 pixels and then separated based on ligand condition. This particle stack was then subjected to classification via several rounds of heterogenous refinement without imposed symmetry. For these refinements, the input classes were four “good” 3D classes generated via *ab initio* reconstruction using manually selected 2D classes and eight “decoy” classes which were generated from random sampling of a small number of particles without alignment. The resultant particle stack of 843,201 particles was then subjected to reference-based motion correction^49^ in Relion version 3.1.3. All further cryo-EM analysis was done in CryoSPARC version 3.2.

A consensus map was then generated using non-uniform refinement with C4 symmetry imposed^50^. During this non-uniform refinement step, spherical aberration, tetrafoil, and anisotropic magnification terms were corrected, and CTF parameters were further optimized. This particle stack was then subjected to three-dimensional classification with 24 classes to classify conformational heterogeneity in the data. The particle stack separated into five main three-dimensional classes: resting (86,623 particles), preactivated (81,483 particles), preactivated-transition (379,443 particles), and inhibited states (178,665 particles), as well as one low resolution “junk” class (116,987 particles).

The resting, preactivated, and inhibited state particle stacks were split by ligand condition and symmetry expanded. For the resting and preactivated particle stacks, focused refinements were then used to improve density for specific domains. Masks were generated for a single hIP_3_R2 protomer, the entire CD, the TMD + JD, ARM3 + CLD, ARM2, and BTF1 + BTF2 + ARM1 in ChimeraX^51,52^. These masks were imported into CryoSPARC and used to focus alignment during local refinement for each resting and preactivated particle stack. For each state and each ligand condition, maps generated using local refinement were used to generate composite maps using Phenix version 1.21.2-5419 Combine Focused Maps^53^. Composite maps were density modified using Phenix Resolve Cryo-EM^54^ and atomic models were built against density modified maps using Coot^55^ version 0.9.2. For the inhibited state, focused refinement was performed only on a single hIP_3_R2 protomer for each ligand condition.

For the preactivated-transition particle stack, three-dimensional variability analysis^56^ was used to separate the data into asymmetric states. Three-dimensional variability analysis of the preactivated-transition particle stack yielded four asymmetric classes: preactivated transition class one (three resting protomers to one preactivated protomer), preactivated transition class two (two resting and two preactivated protomers, opposite each other), preactivated transition class three (two resting and two preactivated protomers, adjacent to each other), and preactivated transition class four (one resting protomer and three preactivated protomers).

### Molecular dynamics simulations

All molecular dynamics simulations were conducted using OpenMM version 7.7.0^57^. The simulation constructs were generated from the experimentally determined apo hIP_3_R2 resting state, the ATP-bound hIP_3_R2 resting state, or the cAMP-bound resting state. The simulation constructs were generated from residues 1909 to 2216 and 2549 to 2636 for each state. For other ligand-bound constructs, the ATP molecule in the ATP-bound construct was replaced with either ADP, AMP, adenosine, or guanosine. Zn^2+^ was kept in all constructs unless otherwise noted. Histidine protonation states were set according to the PropKa prediction at pH 7 within the CHARMMGUI solution builder^58,59,60^. To model a physically realistic Zn^2+^-finger fold^61,62^, Cys2562 and Cys2565 were modeled as cysteine thiolates and His2582 and His2587 were modeled as epsilon-protonated histidine tautomers. Simulations performed without Zn^2+^ showed a destabilized JD even with ATP bound (Supplemental Figure 13A-B) compared to simulations with Zn^2+^. Zn^2+^ was therefore included in all subsequent simulation constructs. The four termini of each reduced construct were patched as standard N and C termini within CHARMMGUI. To limit flexibility at the four termini, harmonic restraints (20 kJ/mol/nm^2^) were applied to the backbone atoms of residues at the four termini of each construct (residues 1909-1915, 2214-2216, 2549-2551, and 2634-2636) during simulation. For constructs with residue substitutions, mutations were made with the CHARMMGUI solution builder. Constructs were solvated in explicit waters based on the TIP3P water model^63^ with 416 mM neutralizing NaCl to match the conditions used for vitrification using the CHARMMGUI solution builder. Further simulation inputs were generated for simulation using OpenMM using the CHARMMGUI solution builder. Following preparation, constructions were energy minimized with an energy tolerance of 100 kJ/mol and were equilibrated for 5 ns with positional restraints on protein sidechains (40 kJ/mol/nm^2^) and backbone (400 kJ/mol/nm^2^). Equilibration was conducted using a Langevin Integrator in the NPT ensemble^64^. Constructs were then released from restraints and simulated for 500 ns in 10 replicates per construct, with the first 100 ns of each replicate discarded as further equilibration. All simulations were performed using the CHARMM36m force field^65^. Systems were simulated in the NPT ensemble at 303.15 K using a LFMiddle Langevin Integrator^66^ (equivalent to BAOAB with leapfrog-like splitting^67^) with a 4 fs timestep and a collision rate of 1 ps^-1^. Pressure was maintained at 1 atm using a molecular-scaling Monte Carlo barostat. Long range interactions were modeled using the particle mesh Ewald method. Hydrogen masses were repartitioned to 4 amu.

### Analysis of MD trajectories

All trajectories were loaded using MDTraj^68^ version 1.9.7 and numerical calculations were carried out using Numpy^69^ version 1.21.6. To identify regions in the JD which have the strongest ligand-dependent dynamicity, we first calculated the per-residue backbone atom root mean square fluctuation (RMSF) for all combined equilibrated ligand-free and ATP-bound trajectories. We next calculated the absolute difference between the RMSF values in the two conditions. For each residue *i*, the absolute difference between RMSF values was computed as

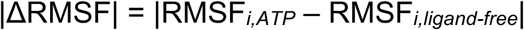

Structured regions in the JD with the highest |ΔRMSF| were selected as candidates for defining the collective variable (CV) that we focused further analysis upon. Specifically, the helices α103 on JD-A and α108 on JD-B were chosen based on the high |ΔRMSF| of residues in these helices. The distance between the backbone atoms of α103 and α108 was chosen as a CV for further analyses.

To quantify the relative motion of α103 and α108, the Euclidean distance between the centers of mass of residues 2176-2183 and 2579-2585 was computed for every trajectory frame. To account for correlation between successive trajectory frames, trajectories were subsampled based on the statistical inefficiency of the distance time series using Pymbar^70^ version 4.0.1. Mean, standard error, and standard deviation of distances are reported from subsampled distributions. To quantify ligand stability in ligand-bound simulations, the Euclidean distance of the center of mass of the bound ligand to the α-carbon of Cys2562 was computed for every trajectory frame. To quantify the interaction frequency of ATP, ADP, AMP, and cAMP with residues in the JD, all pairwise atom-atom distances were computed between bound ligand and protein atoms across all simulation frames. A ligand-residue interaction was defined as occurring in a given frame if any ligand atom was within 3 Å of any atom of a residue.

### Adherent cell culture

HEK293T cells with a stable triple knockout of all three human IP_3_R isoforms^71^ (IP_3_R-null) were acquired from Kerafast. IP_3_R-null cells were maintained in 150 mm cell-culture treated plates (Corning) in DMEM GlutaMax (Thermo Fisher) media supplemented with 10% FBS at 37 C with 5% CO2. For imaging, IP_3_R-null cells at 70-80% confluency were subcultured at a 1:4 ratio and plated on 35 mm plates coated with poly-D-lysine (Fluorodish). IP_3_R-null cells were detached from plates for subculturing by incubation with 0.25% trypsin for 1 min at room temperature. Trypsin was inhibited by subsequent washing with DMEM GlutaMax supplemented with 10% FBS. IP_3_R-null cells were transduced with either wild-type or mutant hIP_3_R2 24 h following subculturing using P3 baculovirus.

### Evaluation of hIP_3_R2 mutant expression and tetrameric assembly

IP_3_R-null cells expressing either wild-type or mutant hIP_3_R2 constructs for 48 h in 6-well 35 mm wells (STEMCELL Technologies) were washed with PBS without Ca^2+^ or Mg^2+^ and removed from plates via trypsinization and subsequent washing with growth media. Cells were pelleted by centrifugation at 2000 RPM for 5 min. Cells were again washed by resuspension in PBS without Ca^2+^ or Mg^2+^, pelleted by centrifugation at 2000 RPM for 5 min, and flash frozen. Cell pellets (approximately 2 × 10^5^ cells) were lysed in 200 µL of a buffer containing 2% lauryl maltose neopentyl glycol (LMNG), 0.2% CHAPS, 300 mM NaCl, 20 mM HEPES pH 7.4, 1 mM PMSF, 1 µg/mL leupeptin, 1 µg/mL aprotinin, 0.5 mM AEBSF, 1 µg/mL pepstatin A, 1 mM benzamidine, and a few flakes of deoxyribonuclease I by constant inversion at 4°C for 1.5 h. Cell lysates were centrifuged at 16,000 ×g for 45 min. Fluorescence size exclusion chromatography was performed on 100 µL of the resulting supernatant for each sample on a Superose 6 Increase column in a buffer containing 0.06% digitonin, 2 mM dithiothreitol, 450 mM NaCl, and 50 mM Tris-HCl pH 8. eGFP fluorescence was monitored by excitation at 488 nm and emission at 510 nm. Expression and folding were evaluated by comparing elution volumes and peak heights between chromatograms resulting from wild-type or mutant hIP_3_R2 runs. Fluorescence size exclusion chromatography was done for each construct in triplicate with similar results between runs.

### Ca^2+^ imaging, processing, and data analysis

IP_3_R-null cells expressing either wild-type or mutant hIP_3_R2 constructs for 48 h were washed with Live Cell Imaging Solution (Invitrogen, containing 140 mM NaCl, 2.5 mM KCl, 1.8 mM CaCl2, 1.0 mM MgCl2, 20 mM HEPES pH 7.4) and incubated with 5 µM Calbryte 590 AM (AAT Bioquest) dissolved in Live Cell Imaging Solution for 1 h at 37 C 5% CO2. Calbryte 590 AM loaded cells were equilibrated at room temperature for 5 min prior to imaging and received intracellular IP_3_ stimulation by constant perfusion of 100 µM carbachol dissolved in the Live Cell Imaging Solution. Movies of cells responding to carbachol treatment were collected at 20x with a LD Plan-Neofluar 20X/0.4 Korr M27 objective for 10 minutes at 3×3 binning with an exposure time of 250 ms on a Zeiss Axio observer D1 inverted phase-contrast fluorescence microscope with a Axiocam 506 Mono camera (Zeiss). Calbryte 590 imaging was done by exciting the sample at 570 nm and monitoring emission at 588 nm using the X-Cite Series 120Q illumination system and the Zeiss filter set 38 HE.

Movies were processed using ImageJ2^72^ version 2.14.0/1.54f and MathWorks MATLAB R2022a version 9.12.0.1974300. Movie stacks were background subtracted with a 50-pixel rolling ball radius in ImageJ2. A maximum intensity projection was used to generate a difference of gaussian which was subsequently used to segment cells using MATLAB’s Image Processing Toolbox. Time series fluorescence traces were extracted from segmented cells and smoothed using a Savitzky-Golay filter of polynomial order 2. Traces were normalized by Z-score and further baseline corrected using the MATLAB function Baseline fit^73^. Ca^2+^ peaks were detected automatically using MATLAB and then manually inspected for confirmation. Traces with ambiguous or artifactual signals were not used for further data analysis. Data reported are from 4 replicate plates with 2 each being from different days. The mean and 95% confidence interval for signals was calculated using NumPy. Mean proportion and standard deviation of Ca^2+^ response for each construct was calculated using NumPy. Statistical testing on pairs of mean proportions between constructs was done using the Mann-Whitney U-test implemented in SciPy^74^.

## Acknowledgements

We thank Jamie Cheong and Neeraj Harikrishna Girija Paramasivam at the Memorial Sloan Kettering Cancer Center High Performance Computing group for assistance with data generation and processing, Lucie Delemotte for assistance with MD simulation analyses, Melinda Diver for access to laboratory space and the members of the Hite, Chodera, Diver, and Delemotte laboratories for helpful comments on the paper. This study was supported in part by R01-GM141553 (R.K.H.), R35-GM156616 (R.K.H.), NCI-F31-243235 (N.P.), R35-GM152017 (J.D.C), and NIH National Cancer Institute Cancer Center Support Grant P30-CA008748 (R.K.H., J.D.C.). V.B. is supported by the Sawyers Fellowship from Memorial Sloan Kettering Cancer Center, NIGMS-5R25GM130494, and T32-GM132081. A portion of this research was supported by NIH grant R24GM154185 and performed at the Pacific Northwest Center for Cryo-EM (PNCC) with assistance from Omar Davulcu.

## Data availability

Cryo-EM maps were deposited to the EM data bank under accession codes: EMD-74147 (hIP_3_R2 resting state without ligands bound), EMD-74148 (hIP_3_R2 resting state without ligands bound, focus refinement of a single protomer), EMD-74160 (hIP_3_R2 resting state without ligands bound, focus refinement of the cytoplasmic domain), EMD-74159 (hIP_3_R2 resting state without ligands bound, focus refinement of the juxtamembrane domain plus the transmembrane domain), EMD-74161 (hIP_3_R2 resting state without ligands bound, focus refinement of the armadillo repeat 3 domain plus the central linker domain), EMD-74162 (hIP_3_R2 resting state without ligands bound, focus refinement of the armadillo repeat 2), EMD-74163 (hIP_3_R2 resting state without ligands bound, focus refinement of beta trefoil domain 1, beta trefoil domain 2, and armadillo repeat domain 1), EMD-75170 (hIP_3_R2 resting state with ATP bound), EMD-74164 (hIP_3_R2 resting state with ATP bound, focus refinement of a single protomer), EMD-74166 (hIP_3_R2 resting state with ATP bound, focus refinement of the cytoplasmic domain), EMD-74170 (hIP_3_R2 resting state with ATP bound, focus refinement of the juxtamembrane domain and the transmembrane domain), EMD-74171 (hIP_3_R2 resting state with ATP bound, focus refinement of the armadillo repeat 3 and the central linker domain), EMD-74168 (hIP_3_R2 resting state with ATP bound, focus refinement of the armadillo repeat 2), EMD-74169 (hIP_3_R2 resting state with ATP bound, focus refinement of the beta trefoil domain 1, beta trefoil domain 2, and armadillo repeat 1), EMD-75036 (hIP_3_R2 resting state with cAMP bound), EMD-74172 (hIP_3_R2 resting state with cAMP bound, focus refinement of a single protomer), EMD-74173 (hIP_3_R2 resting state with cAMP bound, focus refinement of the cytoplasmic domain), EMD-74174 (hIP_3_R2 resting state with cAMP bound, focus refinement of the juxtamembrane domain and the transmembrane domain), EMD-74175 (hIP_3_R2 resting state with cAMP bound, focus refinement of the armadillo repeat 3 and central linker domain), EMD-74176 (hIP_3_R2 resting state with cAMP bound, focus refinement of the armadillo repeat 2), EMD-74177 (hIP_3_R2 resting state with cAMP bound, focus refinement of the beta trefoil domain 1, beta trefoil domain 2, and armadillo repeat 1), EMD-75035 (hIP_3_R2 preactivated state with IP_3_ bound), EMD-74178 (hIP_3_R2 preactivated state with IP_3_ bound, focus refinement of a single protomer), EMD-74179 (hIP_3_R2 preactivated state with IP_3_ bound, focus refinement of the cytoplasmic domain), EMD-74180 (hIP_3_R2 preactivated state with IP_3_ bound, focus refinement of the juxtamembrane domain and transmembrane domain), EMD-74181 (hIP_3_R2 preactivated state with IP_3_ bound, focus refinement of the armadillo repeat 3 and central linker domain), EMD-74182 (hIP_3_R2 preactivated state with IP_3_ bound, focus refinement of the armadillo repeat 2), EMD-74183 (hIP_3_R2 preactivated state with IP_3_ bound, focus refinement of the beta trefoil domain 1, beta trefoil domain 2, and armadillo repeat 1), EMD-74184 (hIP_3_R2 inhibited state without ligands bound, focus refinement of a single protomer), EMD-74188 (hIP_3_R2 inhibited state with ATP bound, focus refinement of a single protomer), EMD-74185 (hIP_3_R2 inhibited state with cAMP bound, focus refinement of a single protomer), EMD-74186 (hIP_3_R2 inhibited state with IP_3_ bound, focus refinement of a single protomer), EMD-74187 (hIP_3_R2 preactivated transition state with three resting protomers and one preactivated protomer without ligands bound), EMD-74189 (hIP_3_R2 preactivated transition state with three resting protomers and one preactivated protomer with ATP bound), EMD-74191 (hIP_3_R2 preactivated transition state with three resting protomers and one preactivated protomer with cAMP bound), EMD-74192 (hIP_3_R2 preactivated transition state with three resting protomers and one preactivated protomer with IP_3_ bound), EMD-74211 (hIP_3_R2 preactivated transition state with two opposing protomers in the resting state and two opposing protomers in the preactivated state without ligands bound), EMD-74242 (hIP_3_R2 preactivated transition state with two opposing protomers in the resting state and two opposing protomers in the preactivated state with ATP bound), EMD-74210 (hIP_3_R2 preactivated transition state with two opposing protomers in the resting state and two opposing protomers in the preactivated state with cAMP bound), EMD-74212 (hIP_3_R2 preactivated transition state with two opposing protomers in the resting state and two opposing protomers in the preactivated state with IP_3_ bound), EMD-74219 (hIP_3_R2 preactivated transition state with two adjacent protomers in the resting state and two adjacent protomers in the preactivated state without ligands bound), EMD-74214 (hIP_3_R2 preactivated transition state with two adjacent protomers in the resting state and two adjacent protomers in the preactivated state with ATP bound), EMD-74216 (hIP_3_R2 preactivated transition state with two adjacent protomers in the resting state and two adjacent protomers in the preactivated state with cAMP bound), EMD-74217 (hIP_3_R2 preactivated transition state with two adjacent protomers in the resting state and two adjacent protomers in the preactivated state with IP_3_ bound), EMD-74234 (hIP_3_R2 preactivated transition state with one resting protomer and three preactivated protomers without ligands bound), EMD-74221 (hIP_3_R2 preactivated transition state with one resting protomer and three preactivated protomers with ATP bound), EMD-74222 (hIP_3_R2 preactivated transition state with one resting protomer and three preactivated protomers with cAMP bound), and EMD-74233 (hIP_3_R2 preactivated transition state with one resting protomer and three preactivated protomers with IP_3_ bound). Atomic models were deposited to the PDB under accession codes: 9ZFP (hIP_3_R2 resting state without ligands bound), 10AV (hIP_3_R2 resting state with ATP bound), 10AU (hIP_3_R2 resting state without ligands bound), 10AV (hIP_3_R2 resting state with cAMP bound), and 10AU (hIP_3_R2 preactivated state with IP_3_ bound). Molecular dynamics simulation trajectories and associated analyses can be replicated via https://github.com/choderalab/ip3r_md.

## Disclaimer

The content is solely the responsibility of the authors and does not necessarily represent the official views of the National Institutes of Health. Any opinions, findings, and conclusions or recommendations expressed in this material are those of the authors and do not necessarily reflect the views of the National Science Foundation.

This manuscript is the result of funding in whole or in part by the National Institutes of Health (NIH). It is subject to the NIH Public Access Policy. Through acceptance of this federal funding, NIH has been given a right to make this manuscript publicly available in PubMed Central upon the Official Date of Publication, as defined by NIH.

