## Supplementary Data for "Structural dynamics of adenine nucleotide potentiation of the human type 2 IP_3_ receptor"

### Supplementary Figures

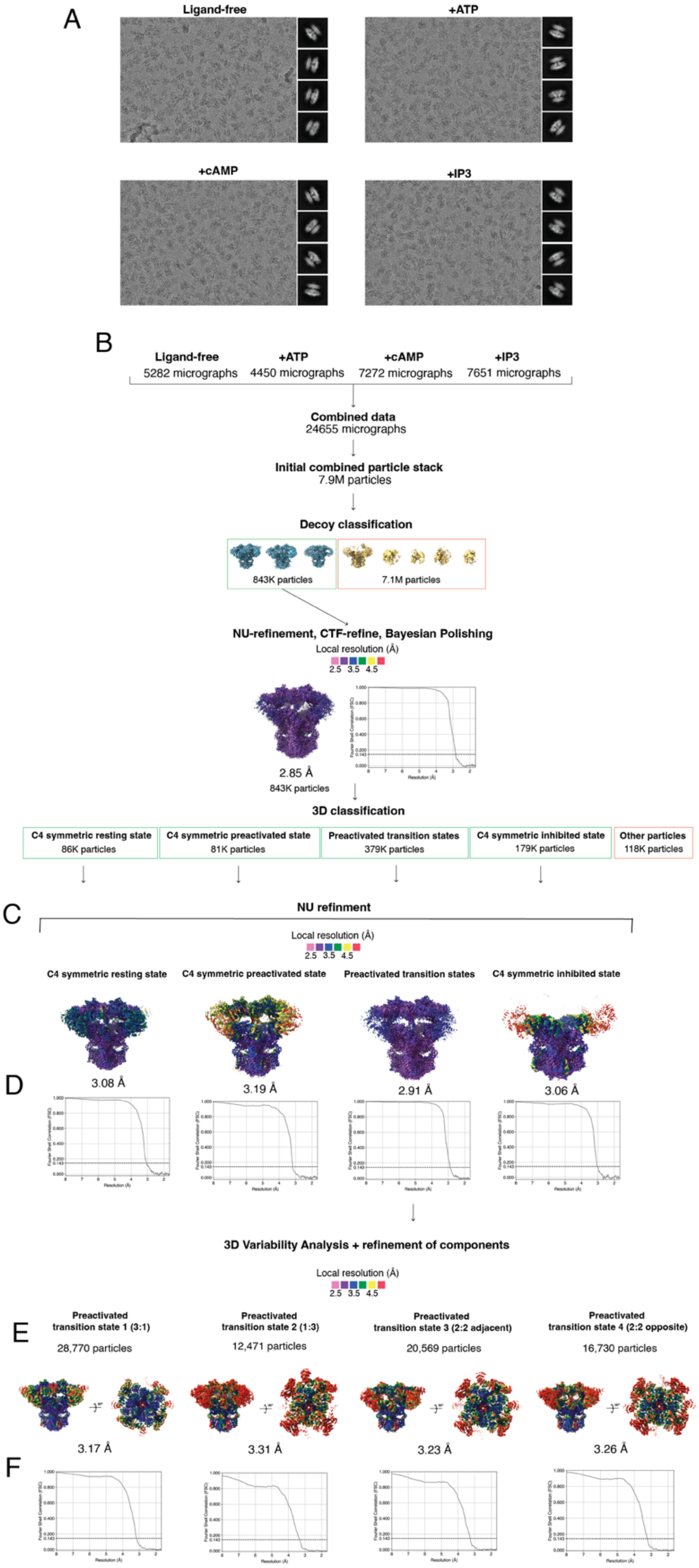
A
Ligand-free
+ATP
+cAMP
+IP3
B
Ligand-free
5282 micrographs
+ATP
4450 micrographs
+cAMP
7272 micrographs
+IP3
7651 micrographs
Combined data
24655 micrographs
Initial combined particle stack
7.9M particles
Decoy classification
843K particles
7.1M particles
NU-refinement, CTF-refine, Bayesian Polishing
Local resolution (Å)
2.5 3.5 4.5
2.85 Å
843K particles
3D classification
C4 symmetric resting state
86K particles
C4 symmetric preactivated state
81K particles
Preactivated transition states
379K particles
C4 symmetric inhibited state
179K particles
Other particles
118K particles
C
NU refinment
Local resolution (Å)
2.5 3.5 4.5
C4 symmetric resting state
C4 symmetric preactivated state
Preactivated transition states
C4 symmetric inhibited state
D
3.08 Å
3.19 Å
2.91 Å
3.06 Å
3D Variability Analysis + refinement of components
Local resolution (Å)
2.5 3.5 4.5
E
Preactivated
transition state 1 (3:1)
28,770 particles
Preactivated
transition state 2 (1:3)
12,471 particles
Preactivated
transition state 3 (2:2 adjacent)
20,569 particles
Preactivated
transition state 4 (2:2 opposite)
16,730 particles
3.17 Å
3.31 Å
3.23 Å
3.26 Å
F

**Supplemental figure 1: Cryo-EM processing and classification workflow.**

**A:** Representative micrographs and 2D class averages for the ligand-free, +ATP, +cAMP, and +$IP_3$ datasets.
**B:** Particle classification and refinement workflow for pooled micrographs. Total number of micrographs are shown per ligand condition. Total particle counts are shown for each classification and refinement step.
**C:** Consensus reconstructions of merged resting, preactivated, transition states and inhibited states, colored by local resolution.
**D:** Half-map FSC plots for consensus reconstructions of merged resting, preactivated, transition states and inhibited states.
**E:** Consensus reconstructions of preactivated transition states, colored by local resolution
**F:** Half-map FSC plots per domain local refinement for consensus reconstructions of preactivated transition states.

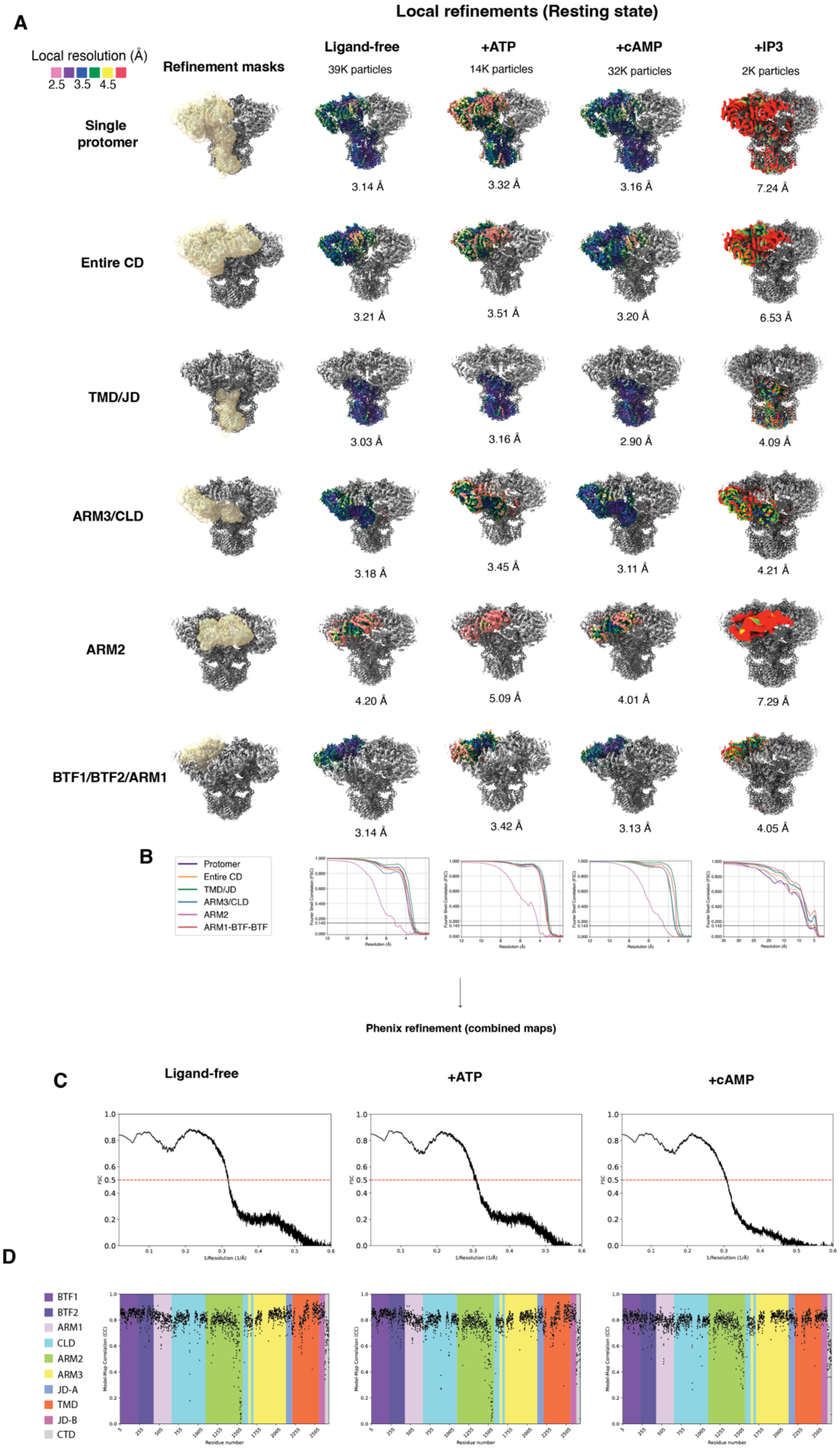
A
Local refinements (Resting state)
Local resolution (Å)
2.5 3.5 4.5
Refinement masks
Ligand-free
39K particles
+ATP
14K particles
+cAMP
32K particles
+IP3
2K particles
Single protomer
3.14 Å
3.32 Å
3.16 Å
7.24 Å
Entire CD
3.21 Å
3.51 Å
3.20 Å
6.53 Å
TMD/JD
3.03 Å
3.16 Å
2.90 Å
4.09 Å
ARM3/CLD
3.18 Å
3.45 Å
3.11 Å
4.21 Å
ARM2
4.20 Å
5.09 Å
4.01 Å
7.29 Å
BTF1/BTF2/ARM1
3.14 Å
3.42 Å
3.13 Å
4.05 Å
B
Protomer
Entire CD
TMD/JD
ARM3/CLD
ARM2
ARM1-BTF-BTF
Phenix refinement (combined maps)
C
Ligand-free
+ATP
+cAMP
FSC
1/Resolution (1/Å)
D
BTF1
BTF2
ARM1
CLD
ARM2
ARM3
JD-A
TMD
JD-B
CTD
Model-Map Correlation (CC)
Residue number

**Supplemental figure 2: Processing of focused and composite maps for the hIP$_3$R2 resting state.**

**A:** Schematic of local refinements performed on particles classified as ligand-free, ATP-bound, cAMP-bound, and IP$_3$-bound resting states. Masks used for local refinement targeting are shown overlaid on the composite resting state tetramer in pale yellow. Focused refinement reconstructions are colored by local resolution. Total particle numbers for each ligand condition in the resting state and nominal resolutions (FSC = 0.143) for each local refinement are shown.
**B:** Half-map FSC plots per domain local refinement for each ligand condition
**C:** Map versus model FSC plots for the composite maps for each ligand condition.
**D:** Per-residue cross-correlation (CC) map versus model plots for composite maps for each ligand condition, with each domain colored uniquely.

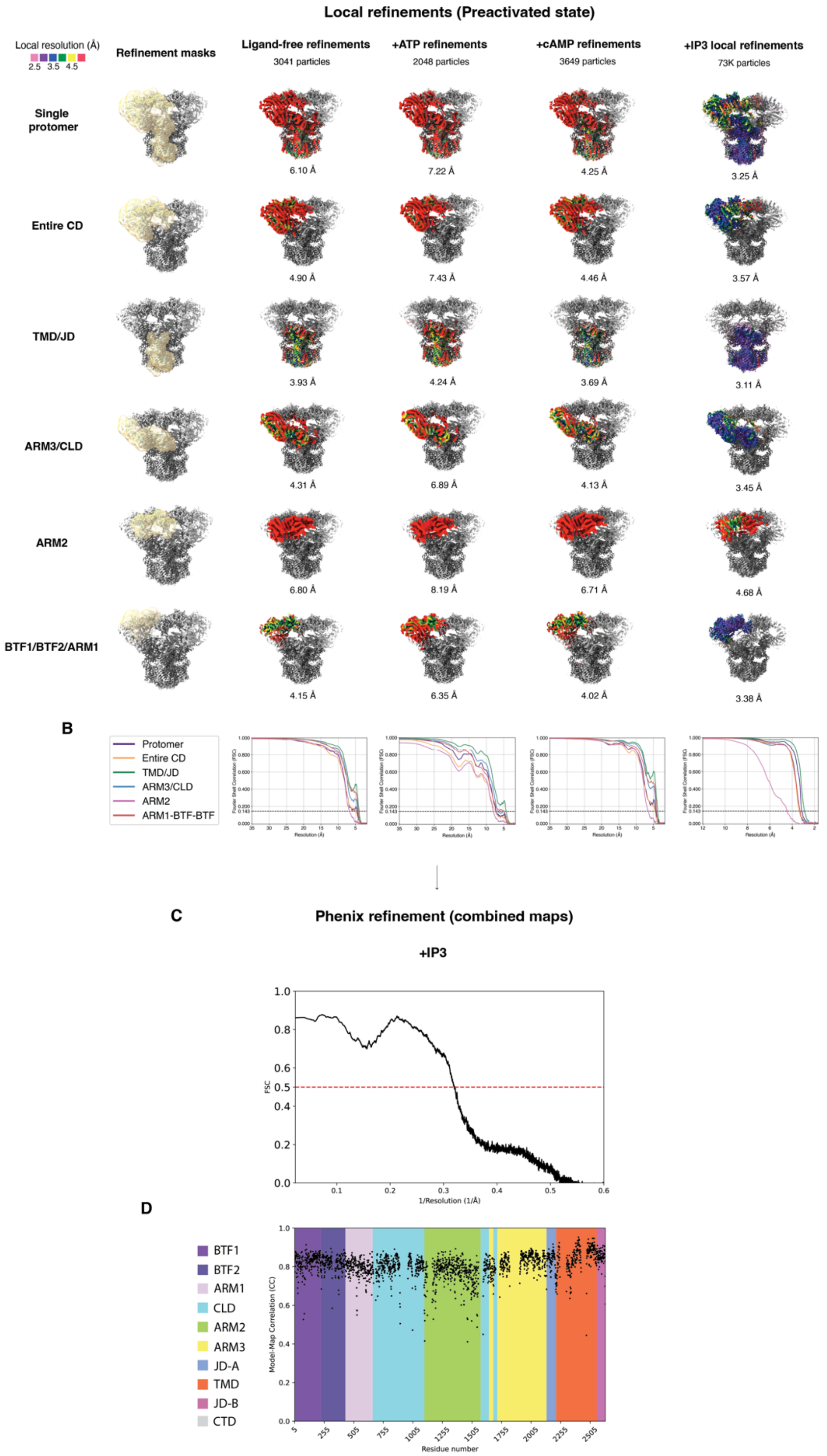
A
Local refinements (Preactivated state)
Local resolution (Å)
2.5 3.5 4.5
Refinement masks
Ligand-free refinements
3041 particles
+ATP refinements
2048 particles
+cAMP refinements
3649 particles
+IP3 local refinements
73K particles
Single protomer
6.10 Å
7.22 Å
4.25 Å
3.25 Å
Entire CD
4.90 Å
7.43 Å
4.46 Å
3.57 Å
TMD/JD
3.93 Å
4.24 Å
3.69 Å
3.11 Å
ARM3/CLD
4.31 Å
6.89 Å
4.13 Å
3.45 Å
ARM2
6.80 Å
8.19 Å
6.71 Å
4.68 Å
BTF1/BTF2/ARM1
4.15 Å
6.35 Å
4.02 Å
3.38 Å
B
Protomer
Entire CD
TMD/JD
ARM3/CLD
ARM2
ARM1-BTF-BTF
Fourier Shell Correlation (FSC)
Resolution (Å)
C
Phenix refinement (combined maps)
+IP3
FSC
1/Resolution (1/Å)
D
BTF1
BTF2
ARM1
CLD
ARM2
ARM3
JD-A
TMD
JD-B
CTD
Model-Map Correlation (CC)
Residue number

**Supplemental figure 3: Processing of focused and composite maps for the $hIP_3R2$ preactivated state.**

**A:** Schematic of local refinements performed on particles classified into the ligand-free, ATP-bound, cAMP-bound, and $IP_3$-bound preactivated states. Masks used for local refinement targeting a shown overlaid on the composite resting state tetramer in pale yellow. Focused refinement reconstructions are colored by local resolution. Total particle numbers for each ligand condition in the resting state and nominal resolutions (FSC = 0.143) for each local refinement are shown.

**B:** Half-map FSC plots per domain local refinement for each ligand condition.

**C:** Map versus model FSC plot for the full length $IP_3$-bound preactivated state model and composite map.

**D:** Per-residue cross-correlation (CC) map versus model plots for the full length $IP_3$-bound preactivated state model and composite map with each domain colored uniquely.

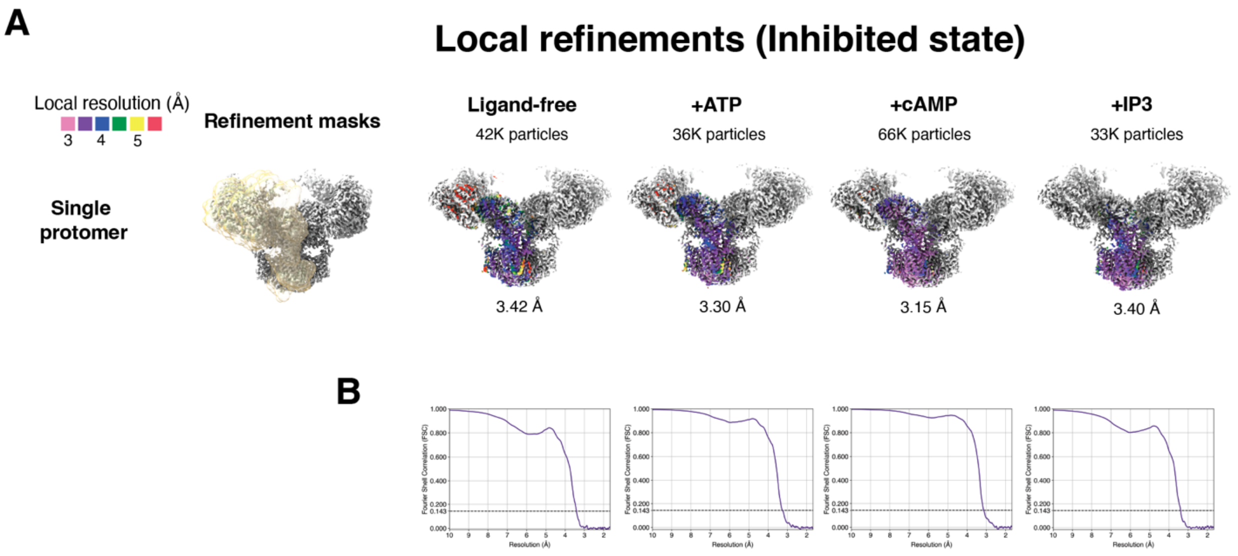

**Supplemental figure 4: Processing of focused and composite maps for the hIP$_3$R2 inhibited state**

**A:** Schematic of local refinements performed on particles classified into the ligand-free, ATP-bound, cAMP-bound, and IP$_3$-bound inhibited states. Masks used for local refinement targeting a shown overlaid on the composite resting state tetramer in pale yellow. Total particle numbers for each ligand condition in the resting state and nominal resolutions (FSC = 0.143) for each local refinement are shown.

**B:** Half-map FSC = 0.143 plots per domain local refinement for each ligand condition.

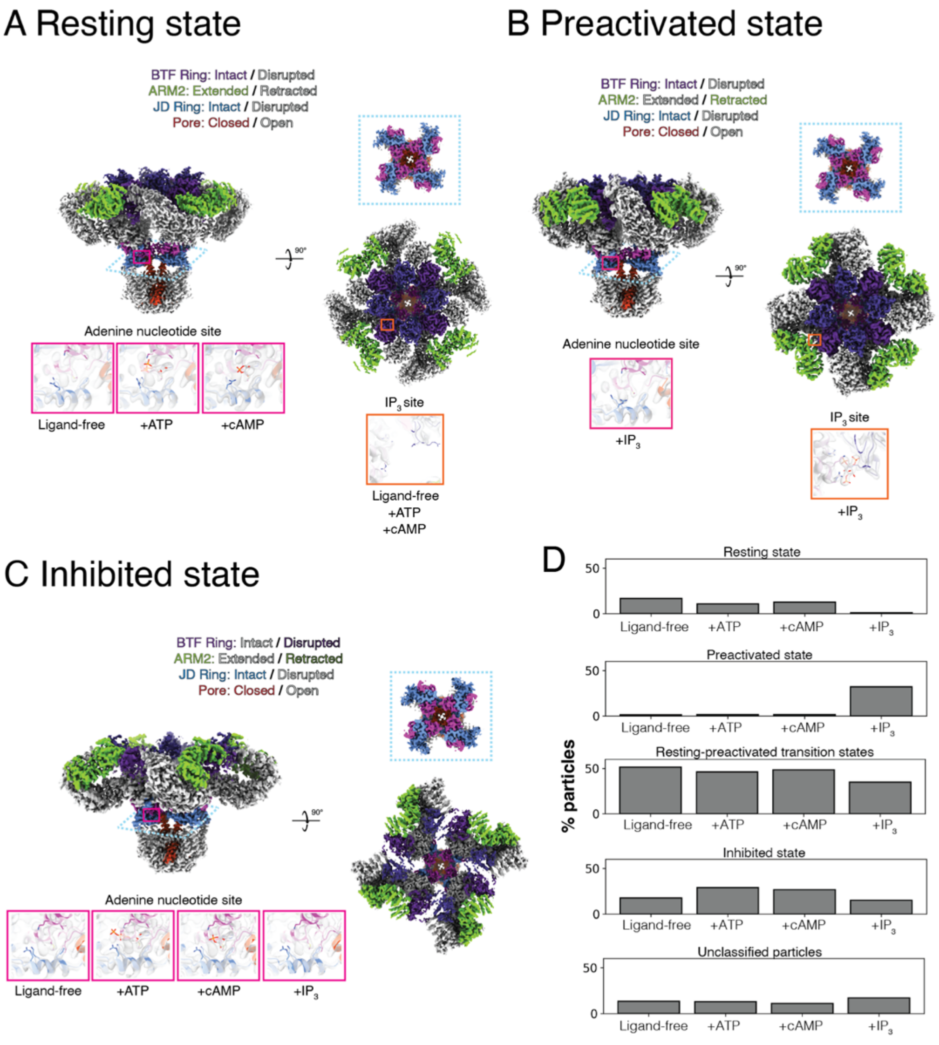

**Supplemental figure 5: C4 symmetric hIP$_3$R2 conformational states.**

**A-C:** Top and side views of tetrameric composite maps of hIP$_3$R2 in resting (A), preactivated (B), and inhibited states (C). Insets show non-protein density in either the IP$_3$- or adenine-nucleotide binding sites. Dashed blue lines correspond to a section through the JD.

**D:** Percent abundance of particles in the resting, preactivated, inhibited, resting-preactivated transition, or unclassified states per ligand condition.

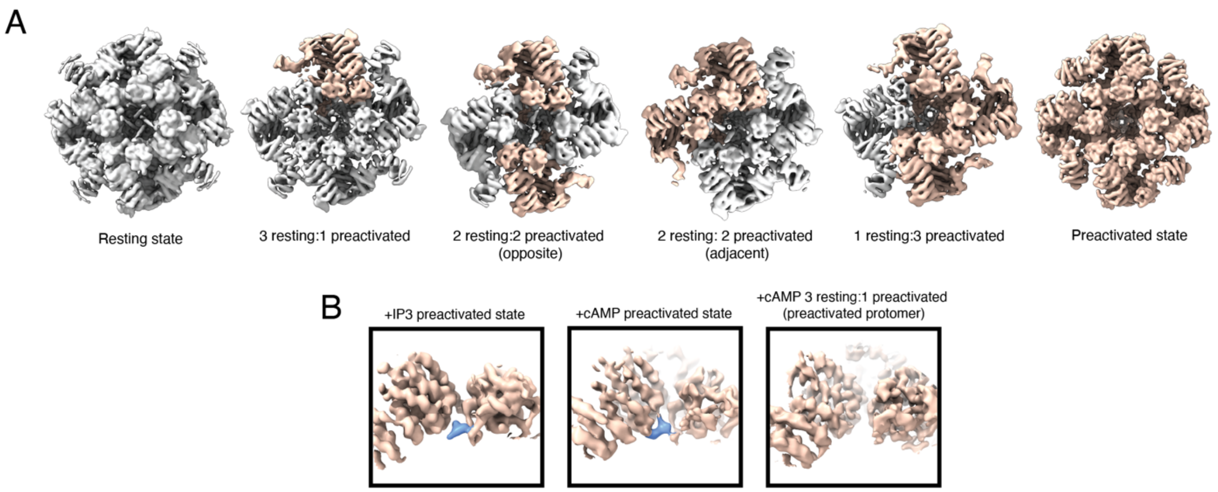

**Supplemental figure 6: Asymmetric h$IP_3$R2 resting-preactivated transition states.**
**A:** Asymmetric resting-preactivated transition states of h$IP_3$R2, viewed from the cytosol. Resting state protomers are colored grey and preactivated state protomers are colored light brown. Maps are low pass filtered to ~5 Å.
**B:** Cryo-EM density maps of the $IP_3$-binding site in either the $IP_3$-bound C4 symmetric preactivated state (low pass filtered to ~4.5 Å), cAMP-bound C4 preactivated state, or the cAMP-bound 3 resting: 1 preactivated state (preactivated protomer displayed). Protein density is shown in light brown and non-protein density is shown in dark blue.

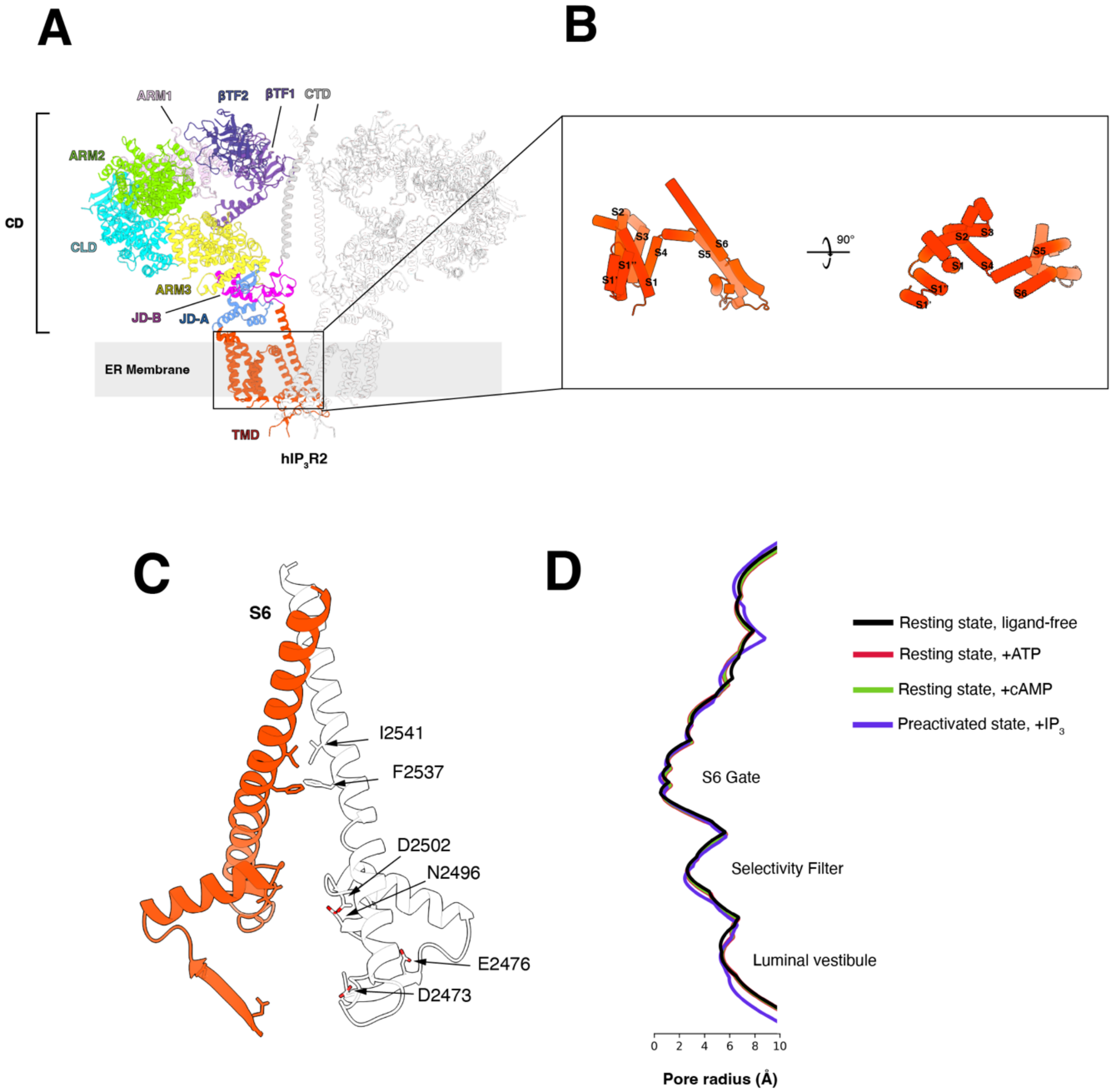

**Supplemental figure 7: Architecture of the hIP$_3$R2 TMD and pore.**

**A:** hIP$_3$R2 in the ligand-free resting state, shown from the side. Color represents protein domain: BTF1 (purple), BTF2 (blue), ARM1 (lavender), ARM2 (green), ARM3 (yellow), CLD (cyan), JD-A (light blue), JD-B (magenta), TMD (red), and CTD (silver). Density for ATP is shown in light yellow. Front and rear protomers are removed for clarity.

**B:** TMD of a single protomer of the ligand-free resting state.

**C:** Structure of the pore-lining S6 helices, with one colored red and the other white. Residues that contribute to pore constrictions are shown as sticks. Front and rear protomers are removed for clarity.

**D:** Pore radius of the ligand-free resting state (black), or bound with ATP (red) or cAMP (green), and the preactivated state bound with IP$_3$ (purple).

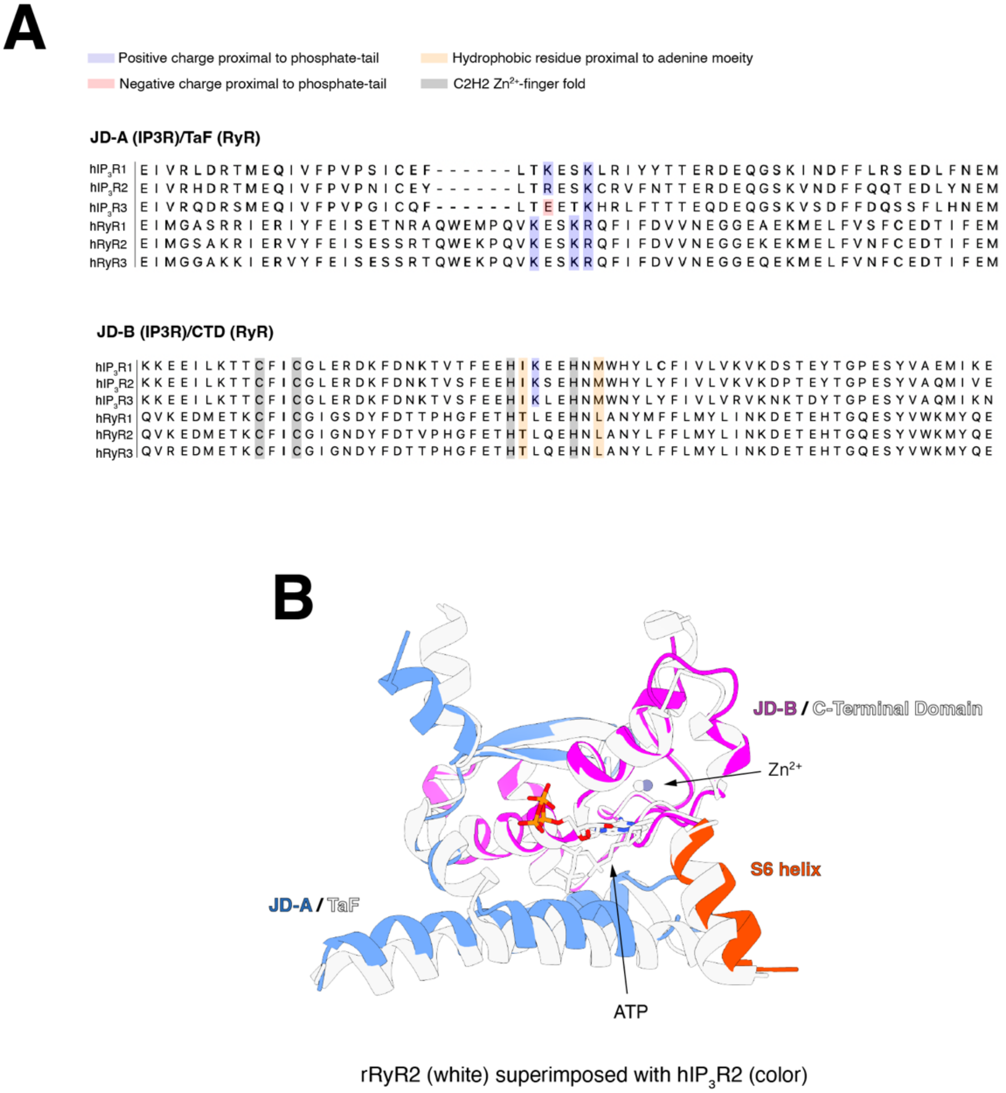

**Supplemental figure 8: The adenine nucleotide binding site and nearby domains are conserved in $IP_3Rs$ and Ryanodine Receptors.**

**A:** Sequence alignment of residues that contribute to adenine nucleotide binding in human $IP_3R$ and Ryanodine Receptors (RyR) isoforms. Regions in the sequence that participate in ATP coordination are highlighted. Positively charged residues are highlighted in blue, negatively charged residues are highlighted in red, hydrophobic residues are highlighted in orange, and residues that form the JD $Zn^{2+}$-finger fold are highlighted in gray.

**B:** Superposition of the ATP-bound resting state of $hIP_3R2$ (shown in color) and the Thumb and Forefingers (TaF) domain and C-Terminal Domain (CTD) of the ATP-bound proteoliposomal primed state of rabbit RyR2 (shown as white, PDB ID: 7M6A), both shown bound with ATP and $Zn^{2+}$.

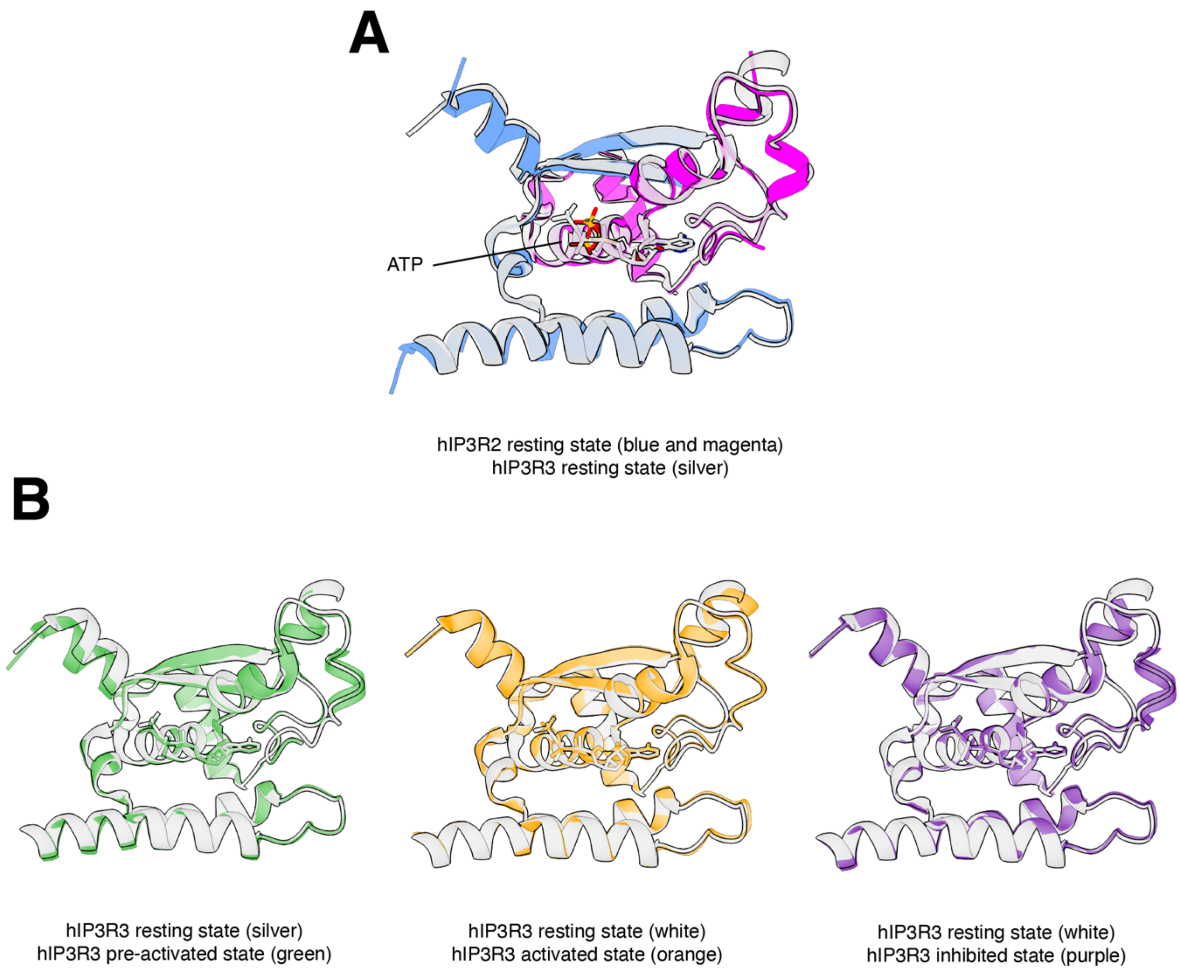

**Supplemental figure 9: JD-A and JD-B move as a rigid body during IP$_3$R activation.**
**A:** Superposition of the JD from the ATP-bound resting states of hIP$_3$R2 (colored by domain) and hIP$_3$R3 (silver).
**B:** Superposition of the JD from hIP$_3$R3 resting state (silver) with the hIP$_3$R3 pre-activated state (left; green), hIP$_3$R3 activated state (center; orange), and hIP$_3$R3 inhibited state (right; purple).

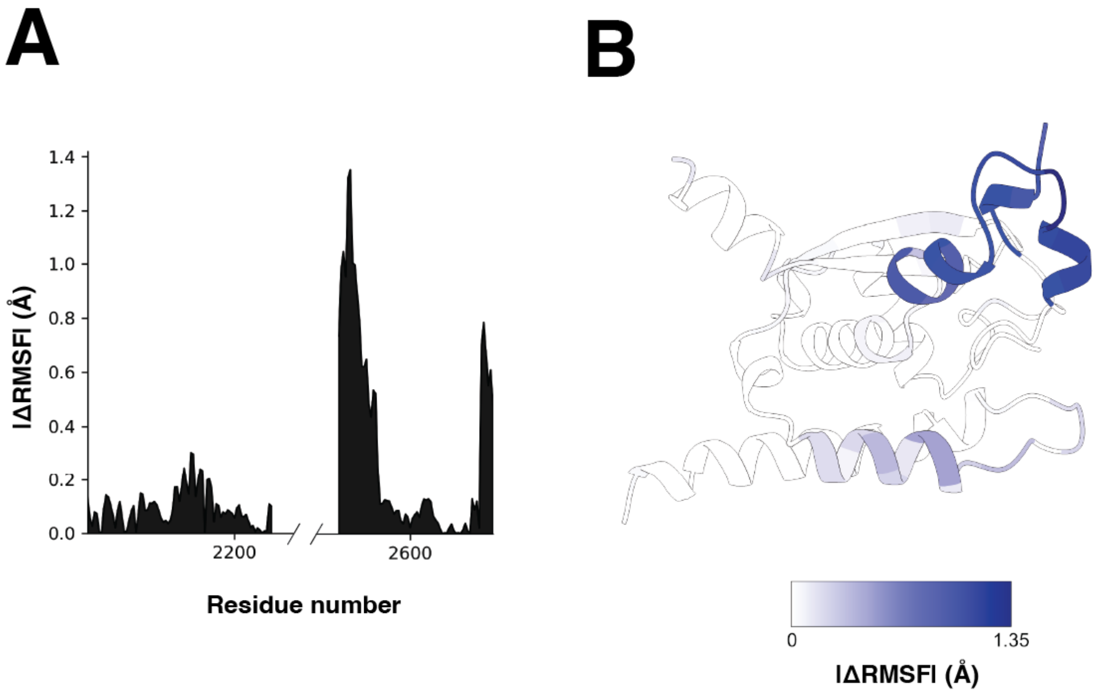

**Supplemental figure 10: Analysis of ligand-dependent root mean square (RMSF) fluctuations in the JD.**
**A:** Per-residue absolute difference in RMSF (|ΔRMSF|) in the JD between ligand-free and ATP-bound trajectories.
**B:** Structure of the JD from the ATP-bound resting state colored by per-residue absolute difference in RMSF.

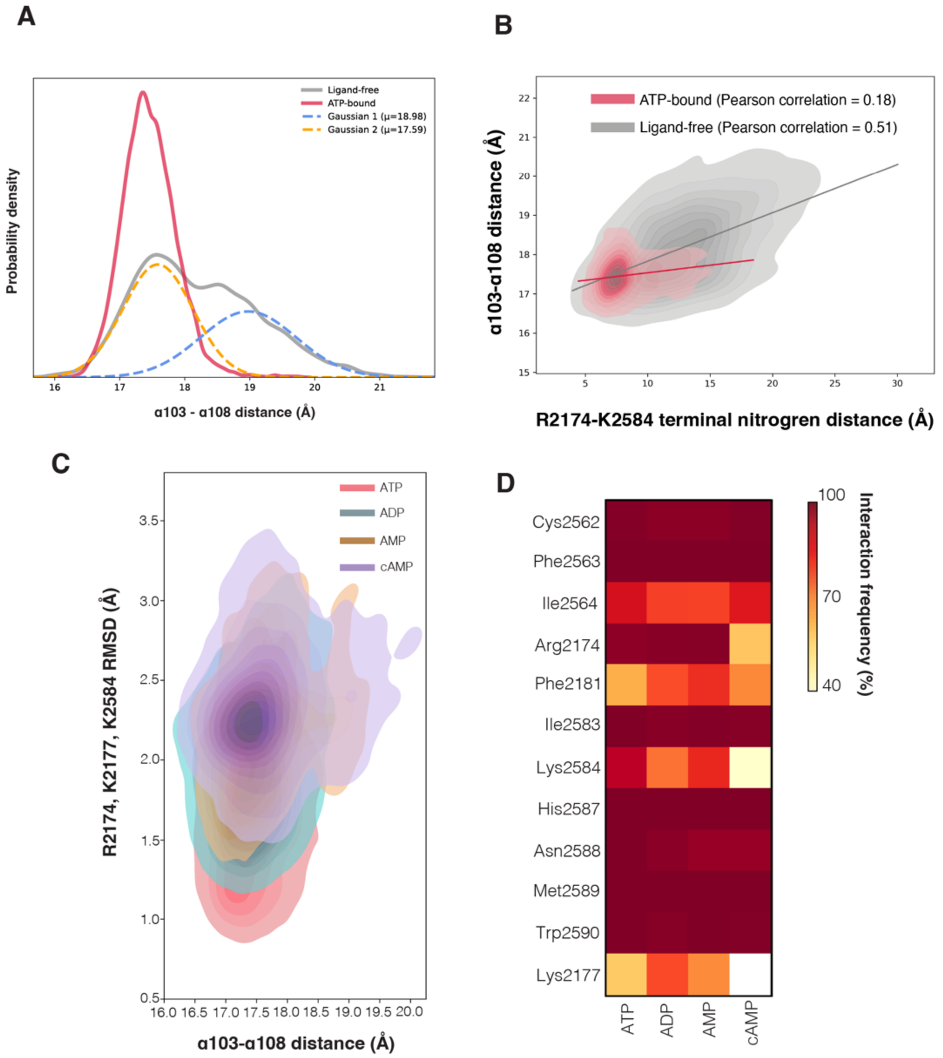

**Supplemental figure 11: Comparative analysis of structural dynamics and residue interaction for ATP-, ADP-, AMP-, and cAMP-bound JD simulation systems.**

**A:** KDEs of ligand-free (gray solid line) and ATP-bound (red solid line) α103/α108 COM distance with two gaussians fit to the ligand-free α103/α108 COM distance distribution (orange and blue dashed lines). Ligand-free and ATP-bound α103/α108 COM distance are the same as presented in figure 2.

**B:** 1D and 2D KDEs of the relationship between the COM distance between all atoms in R2174 and K2584, and the COM distance between the backbone atoms of α103 and α108 in ATP-bound (red) or ligand-free states (gray). Solid lines represent linear regression fits for either measurement. ATP-bound α103/α108 COM distance is the same as presented in figure 2.

**C:** 2D KDE of the relationship between the α103/α108 COM distance and the RMSD of Arg2174, Lys2177, and Lys2584 from ATP-bound, ADP-bound, AMP-bound, or cAMP-bound trajectories. ATP-bound α103/α108 COM distance is the same as presented in figure 2.

**D:** Heat map of residue-nucleotide interactions for the ATP, ADP, AMP, and cAMP simulations.

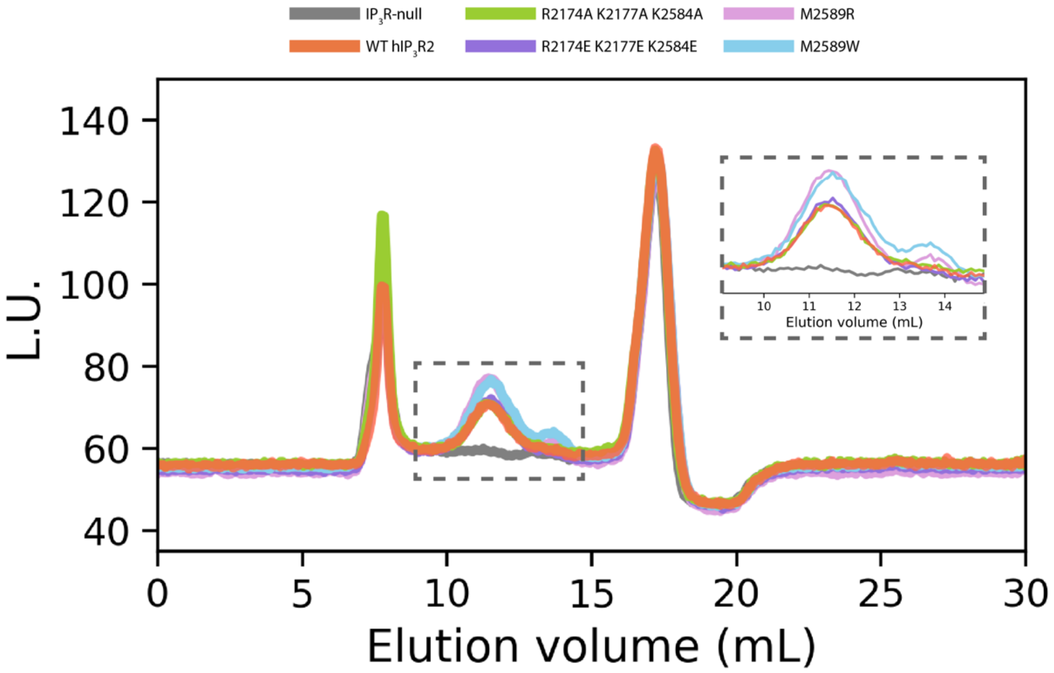

**Supplemental figure 12: Adenine nucleotide binding site mutations minimally impact $hIP_3R2$ expression or tetrameric assembly.**
Fluorescence size exclusion chromatography profiles of the soluble fraction of detergent-solubilized whole cell lysate from $IP_3R$-null cells and $IP_3R$-null cells transduced with wild-type or mutant $hIP_3R2$. Dashed grey box corresponds to inset at top right.

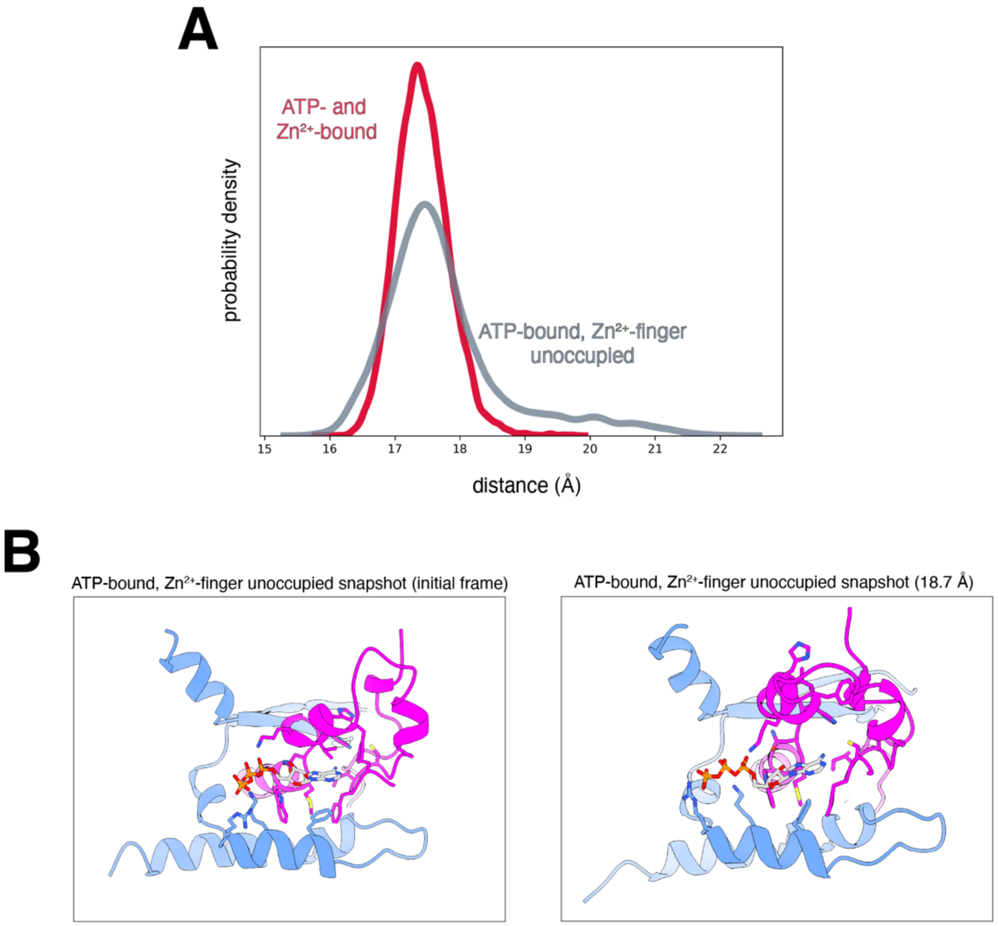

**Supplemental figure 13: $Zn^{2+}$ is required for ATP-dependent JD stability in MD simulations**

**A:** KDE of the COM distance between the backbone atoms of α103 and α108 from the ATP+$Zn^{2+}$-bound (red) and ATP-bound (gray) trajectories.

**B:** Structural snapshots of JD-A and JD-B with ATP bound and without $Zn^{2+}$ illustrating structural flexibility in the absence of $Zn^{2+}$.

### Supplemental table 1: Cryo-EM statistics.

| **Data collection and processing** | #1 hIP3R2 resting state, ligand-free<br>EMD-74147<br>PDB 9ZFP | #2 hIP3R2 resting state, ligand-free, single protomer<br>EMD-74148 | #3 hIP3R2 resting state, ligand-free, cytoplasmic domain<br>EMD-74160 | #4 hIP3R2 resting state, ligand-free, juxtamembrane domain and transmembrane domain<br>EMD-74159 | #5 hIP3R2 resting state, ligand-free, armadillo repeat 3 and central linker domain<br>EMD-74161 | #6 hIP3R2, ligand-free, armadillo repeat 2<br>EMD-74162 | #7 hIP3R2 resting state, ligand-free, beta trefoil 1, beta trefoil 2, and armadillo repeat 1<br>EMD-74163 |
|---|---|---|---|---|---|---|---|
| Detector | Gatan K3 | Gatan K3 | Gatan K3 | Gatan K3 | Gatan K3 | Gatan K3 | Gatan K3 |
| Magnification | 29,000X | 29,000X | 29,000X | 29,000X | 29,000X | 29,000X | 29,000X |
| Voltage (kV) | 300 | 300 | 300 | 300 | 300 | 300 | 300 |
| Electron exposure (e–/Å$^2$) | 66 | 66 | 66 | 66 | 66 | 66 | 66 |
| Defocus range (μm) | -0.1 to -3 | -0.1 to -3 | -0.1 to -3 | -0.1 to -3 | -0.1 to -3 | -0.1 to -3 | -0.1 to -3 |
| Super-resolution pixel size (Å) | 0.413 | 0.413 | 0.413 | 0.413 | 0.413 | 0.413 | 0.413 |
| Final pixel size (Å) | 0.826 | 0.826 | 0.826 | 0.826 | 0.826 | 0.826 | 0.826 |
| Symmetry imposed | C4 | C1 | C1 | C1 | C1 | C1 | C1 |
| Initial particle images (no.) | 7,961,605 | 7,961,605 | 7,961,605 | 7,961,605 | 7,961,605 | 7,961,605 | 7,961,605 |
| Final particle images (no.) | 38,938 | 38,938 | 38,938 | 38,938 | 38,938 | 38,938 | 38,938 |
| Map resolution (Å)<br>FSC threshold | 3.14<br>0.143 | 3.14<br>0.143 | 3.21<br>0.143 | 3.03<br>0.143 | 3.18<br>0.143 | 4.2<br>0.143 | 3.14<br>0.143 |
| **Refinement** | | | | | | | |
| Model resolution (Å)<br>0.5 FSC threshold | 3.16 | | | | | | |
| Model composition<br>Non-hydrogen atoms<br>Protein residues<br>Ligands | <br>72,620<br>2,242<br>1 | | | | | | |
| Mean *B* factors (Å$^2$)<br>Protein<br>Ligand<br>Water | <br>94.48<br>95.79<br> | | | | | | |
| R.m.s. deviations<br>Bond lengths (Å)<br>Bond angles (°) | <br>0.12<br>0.27 | | | | | | |
| Validation<br>MolProbity score | <br>0.455 | | | | | | |

| | |
|---|---|
| Clashscore | 5 |
| Poor rotamers (%) | 0.8 |
| Ramachandran plot | |
| Favored (%) | 99 |
| Allowed (%) | 1 |
| Disallowed (%) | 0 |

| **Data collection and processing** | #8 hIP3R2 resting state, ATP-bound<br>EMD-75170<br>PDB 10HH | #9 hIP3R2 resting state, ATP-bound, single protomer<br>EMD-74164 | #10 hIP3R2 resting state, ATP-bound, cytoplasmic domain<br>EMD-74166. | #11 hIP3R2 resting state, ATP-bound, juxtamembrane domain and transmembrane domain<br>EMD-74170 | #12 hIP3R2 resting state, ATP-bound, armadillo repeat 3 and central linker domain<br>EMD-74171 | #13 hIP3R2 resting state, ATP-bound, armadillo repeat 2<br>EMD-74168 | #14 hIP3R2 resting state, ATP-bound, beta trefoil domain 1, beta trefoil domain 2, armadillo repeat 1<br>EMD-74169 |
|---|---|---|---|---|---|---|---|
| Detector | Gatan K3 | Gatan K3 | Gatan K3 | Gatan K3 | Gatan K3 | Gatan K3 | Gatan K3 |
| Magnification | 29,000X | 29,000X | 29,000X | 29,000X | 29,000X | 29,000X | 29,000X |
| Voltage (kV) | 300 | 300 | 300 | 300 | 300 | 300 | 300 |
| Electron exposure (e–/Å$^2$) | 66 | 66 | 66 | 66 | 66 | 66 | 66 |
| Defocus range (μm) | -0.1 to -3 | -0.1 to -3 | -0.1 to -3 | -0.1 to -3 | -0.1 to -3 | -0.1 to -3 | -0.1 to -3 |
| Super-resolution pixel size (Å) | 0.413 | 0.413 | 0.413 | 0.413 | 0.413 | 0.413 | 0.413 |
| Final pixel size (Å) | 0.826 | 0.826 | 0.826 | 0.826 | 0.826 | 0.826 | 0.826 |
| Symmetry imposed | C4 | C1 | C1 | C1 | C1 | C1 | C1 |
| Initial particle images (no.) | 7,961,605 | 7,961,605 | 7,961,605 | 7,961,605 | 7,961,605 | 7,961,605 | 7,961,605 |
| Final particle images (no.) | 13,638 | 13,638 | 13,638 | 13,638 | 13,638 | 13,638 | 13,638 |
| Map resolution (Å) | 3.32 | 3.32 | 3.51 | 3.16 | 3.45 | 5.09 | 3.42 |
| FSC threshold | 0.143 | 0.143 | 0.143 | 0.143 | 0.143 | 0.143 | 0.143 |
| **Refinement** | | | | | | | |
| Model resolution (Å)<br>0.5 FSC threshold | 3.25 | | | | | | |
| Model composition | | | | | | | |
| Non-hydrogen atoms | 72,760 | | | | | | |
| Protein residues | 2,242 | | | | | | |
| Ligands | 2 | | | | | | |
| Mean *B* factors (Å$^2$) | | | | | | | |
| Protein | 96.58 | | | | | | |
| Ligand | 81.93 | | | | | | |
| Water | | | | | | | |
| R.m.s. deviations | | | | | | | |

| | |
|---|---|
| Bond lengths (Å) | 0.11 |
| Bond angles (°) | 0.27 |
| Validation | |
| MolProbity score | 0.445 |
| Clashscore | 6 |
| Poor rotamers (%) | 1 |
| Ramachandran plot | |
| Favored (%) | 98 |
| Allowed (%) | 2 |
| Disallowed (%) | 0 |

| **Data collection and processing** | #15 hIP3R2 resting state, cAMP-bound<br>EMD-75036<br>PDB 10AV | #16 hIP3R2 resting state, cAMP-bound, protomer<br>EMD-74172 | #17 hIP3R2 resting state, cAMP-bound, cytoplasmic domain<br>EMD-74173 | #18 hIP3R2 resting state, cAMP-bound, juxtamembrane domain and transmembrane domain<br>EMD-74174 | #19 hIP3R2 resting state, cAMP-bound, armadillo repeat 3 and central linker domain<br>EMD-74175 | #20 hIP3R2 resting state, cAMP-bound, armadillo repeat 2<br>EMD-74176 | #21 hIP3R2 resting state, cAMP-bound, beta-trefoil domain 1, beta-trefoil domain 2, and armadillo repeat 1<br>EMD-74177 |
|---|---|---|---|---|---|---|---|
| Detector | Gatan K3 | Gatan K3 | Gatan K3 | Gatan K3 | Gatan K3 | Gatan K3 | Gatan K3 |
| Magnification | 29,000X | 29,000X | 29,000X | 29,000X | 29,000X | 29,000X | 29,000X |
| Voltage (kV) | 300 | 300 | 300 | 300 | 300 | 300 | 300 |
| Electron exposure (e–/Å$^2$) | 66 | 66 | 66 | 66 | 66 | 66 | 66 |
| Defocus range (μm) | -0.1 to -3 | -0.1 to -3 | -0.1 to -3 | -0.1 to -3 | -0.1 to -3 | -0.1 to -3 | -0.1 to -3 |
| Super-resolution pixel size (Å) | 0.413 | 0.413 | 0.413 | 0.413 | 0.413 | 0.413 | 0.413 |
| Final pixel size (Å) | 0.826 | 0.826 | 0.826 | 0.826 | 0.826 | 0.826 | 0.826 |
| Symmetry imposed | C1 | C1 | C1 | C1 | C1 | C1 | C1 |
| Initial particle images (no.) | 7,961,605 | 7,961,605 | 7,961,605 | 7,961,605 | 7,961,605 | 7,961,605 | 7,961,605 |
| Final particle images (no.) | 31,509 | 31,509 | 31,509 | 31,509 | 31,509 | 31,509 | 31,509 |
| Map resolution (Å) | 3.13 | 3.2 | 2.9 | 3.25 | 3.11 | 4.01 | 3.13 |
| FSC threshold | 0.143 | 0.143 | 0.143 | 0.143 | 0.143 | 0.143 | 0.143 |
| **Refinement** | | | | | | | |
| Model resolution (Å)<br>0.5 FSC threshold | 3.27 | | | | | | |
| Model composition | | | | | | | |
| Non-hydrogen atoms | 72,708 | | | | | | |
| Protein residues | 2,242 | | | | | | |
| Ligands | 2 | | | | | | |

| | |
|---|---|
| Mean *B* factors (Å$^2$) | |
| Protein | 101.38 |
| Ligand | 90.35 |
| Water | |
| R.m.s. deviations | |
| Bond lengths (Å) | 0.12 |
| Bond angles (°) | 0.29 |
| Validation | |
| MolProbity score | 0.433 |
| Clashscore | 8 |
| Poor rotamers (%) | 0.9 |
| Ramachandran plot | |
| Favored (%) | 98 |
| Allowed (%) | 2 |
| Disallowed (%) | 0 |

| **Data collection and processing** | #22 hIP3R2, preactivated state, IP3-bound<br>EMD-75035.<br>PDB 10AU | #23 hIP3R2, preactivated state, IP3-bound, protomer<br>EMD-74178 | #24 hIP3R2, preactivated state, IP3-bound, cytoplasmic domain<br>EMD-74179 | #25 hIP3R2, preactivated state, IP3-bound, juxtamembrane domain and transmembrane domain<br>EMD-74180 | #26 hIP3R2, preactivated state, IP3-bound, armadillo repeat 3 and central linker domain<br>EMD-74181 | #27 hIP3R2, preactivated state, IP3-bound, armadillo repeat 2<br>EMD-74182. | #28 hIP3R2, preactivated state, IP3-bound, beta trefoil domain 1, beta trefoil domain 2, and armadillo repeat 1<br>EMD-74183. |
|---|---|---|---|---|---|---|---|
| Detector | Gatan K3 | Gatan K3 | Gatan K3 | Gatan K3 | Gatan K3 | Gatan K3 | Gatan K3 |
| Magnification | 29,000X | 29,000X | 29,000X | 29,000X | 29,000X | 29,000X | 29,000X |
| Voltage (kV) | 300 | 300 | 300 | 300 | 300 | 300 | 300 |
| Electron exposure (e–/Å$^2$) | 66 | 66 | 66 | 66 | 66 | 66 | 66 |
| Defocus range (μm) | -0.1 to -3 | -0.1 to -3 | -0.1 to -3 | -0.1 to -3 | -0.1 to -3 | -0.1 to -3 | -0.1 to -3 |
| Super-resolution pixel size (Å) | 0.413 | 0.413 | 0.413 | 0.413 | 0.413 | 0.413 | 0.413 |
| Final pixel size (Å) | 0.826 | 0.826 | 0.826 | 0.826 | 0.826 | 0.826 | 0.826 |
| Symmetry imposed | C1 | C1 | C1 | C1 | C1 | C1 | C1 |
| Initial particle images (no.) | 7,961,605 | 7,961,605 | 7,961,605 | 7,961,605 | 7,961,605 | 7,961,605 | 7,961,605 |
| Final particle images (no.) | 72,745 | 72,745 | 72,745 | 72,745 | 72,745 | 72,745 | 72,745 |
| Map resolution (Å) | 3.17 | 3.17 | 3.35 | 2.93 | 3.3 | 4.41 | 3.26 |
| FSC threshold | 0.143 | 0.143 | 0.143 | 0.143 | 0.143 | 0.143 | 0.143 |
| **Refinement** | | | | | | | |
| Model resolution (Å) | 3.14 | | | | | | |

| | |
|---|---|
| 0.5 FSC threshold | |
| Model composition<br>Non-hydrogen atoms<br>Protein residues<br>Ligands | <br>70,820<br>2155<br>2 |
| Mean *B* factors (Å$^2$)<br>Protein<br>Ligand<br>Water | <br>91.74<br>92.92<br> |
| R.m.s. deviations<br>Bond lengths (Å)<br>Bond angles (°) | <br>0.2<br>0.4 |
| Validation<br>MolProbity score<br>Clashscore<br>Poor rotamers (%) | <br>0.448<br>7<br>1.2 |
| Ramachandran plot<br>Favored (%)<br>Allowed (%)<br>Disallowed (%) | <br>98<br>2<br>0 |

| **Data collection and processing** | #29 hIP3R2, inhibited state, ligand-free, protomer<br><br>EMD-74184 | #30 hIP3R2, inhibited state, ATP-bound, protomer<br><br>EMD-74188. | #31 hIP3R2, inhibited state, cAMP-bound, protomer<br><br>EMD-74185 | #32 hIP3R2, inhibited state, IP3-bound, protomer<br><br>EMD-74186 | #33 hIP3R2, preactivated transition state, 3 resting to 1 preactivated protomer, ligand-free<br><br>EMD-74187. | #34 hIP3R2, preactivated transition state, 3 resting to 1 preactivated protomer, ATP-bound<br><br>EMD-74242 | #35 hIP3R2, preactivated transition state, 3 resting to 1 preactivated protomer, cAMP-bound<br><br>EMD-74210 |
|---|---|---|---|---|---|---|---|
| Detector | Gatan K3 | Gatan K3 | Gatan K3 | Gatan K3 | Gatan K3 | Gatan K3 | Gatan K3 |
| Magnification | 29,000X | 29,000X | 29,000X | 29,000X | 29,000X | 29,000X | 29,000X |
| Voltage (kV) | 300 | 300 | 300 | 300 | 300 | 300 | 300 |
| Electron exposure (e–/Å$^2$) | 66 | 66 | 66 | 66 | 66 | 66 | 66 |
| Defocus range (μm) | -0.1 to -3 | -0.1 to -3 | -0.1 to -3 | -0.1 to -3 | -0.1 to -3 | -0.1 to -3 | -0.1 to -3 |
| Super-resolution pixel size (Å) | 0.413 | 0.413 | 0.413 | 0.413 | 0.413 | 0.413 | 0.413 |
| Final pixel size (Å) | 0.826 | 0.826 | 0.826 | 0.826 | 0.826 | 0.826 | 0.826 |
| Symmetry imposed | C1 | C1 | C1 | C1 | C1 | C1 | C1 |
| Initial particle images (no.) | 7,961,605 | 7,961,605 | 7,961,605 | 7,961,605 | 7,961,605 | 7,961,605 | 7,961,605 |
| Final particle images (no.) | 41,311 | 37,263 | 66,254 | 34,351 | 11,422 | 5,004 | 11,037 |
| Map resolution (Å)<br>FSC threshold | 3.42<br>0.143 | 3.3<br>0.143 | 3.15<br>0.143 | 3.4<br>0.143 | 3.61<br>0.143 | 3.88<br>0.143 | 3.6<br>0.143 |

| Data collection and processing | #36 hIP3R2, preactivated transition state, 3 resting to 1 preactivated protomer, IP3-bound<br><br>EMD-74212 | #37 hIP3R2, preactivated transition state, 2 opposing resting protomers and 2 opposing preactivated protomers, ligand-free<br><br>EMD-74219 | #38 hIP3R2, preactivated transition state, 2 opposing resting protomers and 2 opposing preactivated protomers, ATP-bound<br><br>EMD-74211 | #39 preactivated transition state, 2 opposing resting protomers and 2 opposing preactivated protomers, cAMP-bound<br><br>EMD-74242 | #40 preactivated transition state, 2 opposing resting protomers and 2 opposing preactivated protomers, IP3-bound<br><br>EMD-74210 | #41 hIP3R2, preactivated transition state, 2 adjacent resting protomers and 2 adjacent preactivated protomers, ligand-free<br><br>EMD-74212 | #42 hIP3R2, preactivated transition state, 2 adjacent resting protomers and 2 adjacent preactivated protomers, ATP-bound<br><br>EMD-74219 |
|---|---|---|---|---|---|---|---|
| Detector | Gatan K3 | Gatan K3 | Gatan K3 | Gatan K3 | Gatan K3 | Gatan K3 | Gatan K3 |
| Magnification | 29,000X | 29,000X | 29,000X | 29,000X | 29,000X | 29,000X | 29,000X |
| Voltage (kV) | 300 | 300 | 300 | 300 | 300 | 300 | 300 |
| Electron exposure (e–/Å$^2$) | 66 | 66 | 66 | 66 | 66 | 66 | 66 |
| Defocus range (μm) | -0.1 to -3 | -0.1 to -3 | -0.1 to -3 | -0.1 to -3 | -0.1 to -3 | -0.1 to -3 | -0.1 to -3 |
| Super-resolution pixel size (Å) | 0.413 | 0.413 | 0.413 | 0.413 | 0.413 | 0.413 | 0.413 |
| Final pixel size (Å) | 0.826 | 0.826 | 0.826 | 0.826 | 0.826 | 0.826 | 0.826 |
| Symmetry imposed | C1 | C1 | C1 | C1 | C1 | C1 | C1 |
| Initial particle images (no.) | 7,961,605 | 7,961,605 | 7,961,605 | 7,961,605 | 7,961,605 | 7,961,605 | 7,961,605 |
| Final particle images (no.) | 1,307 | 4,417 | 2,408 | 4,608 | 5,297 | 6,396 | 3,128 |
| Map resolution (Å) | 6.73 | 4.04 | 4.36 | 3.95 | 4.06 | 3.88 | 4.08 |
| FSC threshold | 0.143 | 0.143 | 0.143 | 0.143 | 0.143 | 0.143 | 0.143 |

| Data collection and processing | #43 hIP3R2, preactivated transition state, 2 adjacent resting protomers and 2 adjacent preactivated protomers, cAMP-bound<br><br>EMD-74214 | #44 hIP3R2, preactivated transition state, 2 adjacent resting protomers and 2 adjacent preactivated protomers, IP3-bound<br><br>EMD-74216 | #45 hIP3R2, preactivated transition state, 1 resting to 3 preactivated protomer, ligand-free<br><br>EMD-74217 | #46 hIP3R2, preactivated transition state, 1 resting to 3 preactivated protomer, ATP-bound<br><br>EMD-74234 | #47 hIP3R2, preactivated transition state, 1 resting to 3 preactivated protomer, cAMP-bound<br><br>EMD-74221 | #48 hIP3R2, preactivated transition state, 1 resting to 3 preactivated protomer, IP3-bound<br><br>EMD-74222 |
|---|---|---|---|---|---|---|

| | | | | | | |
|---|---|---|---|---|---|---|
| Detector | Gatan K3 | Gatan K3 | Gatan K3 | Gatan K3 | Gatan K3 | Gatan K3 |
| Magnification | 29,000X | 29,000X | 29,000X | 29,000X | 29,000X | 29,000X |
| Voltage (kV) | 300 | 300 | 300 | 300 | 300 | 300 |
| Electron exposure (e–/Å$^2$) | 66 | 66 | 66 | 66 | 66 | 66 |
| Defocus range (μm) | -0.1 to -3 | -0.1 to -3 | -0.1 to -3 | -0.1 to -3 | -0.1 to -3 | -0.1 to -3 |
| Super-resolution pixel size (Å) | 0.413 | 0.413 | 0.413 | 0.413 | 0.413 | 0.413 |
| Final pixel size (Å) | 0.826 | 0.826 | 0.826 | 0.826 | 0.826 | 0.826 |
| Symmetry imposed | C1 | C1 | C1 | C1 | C1 | C1 |
| Initial particle images (no.) | 7,961,605 | 7,961,605 | 7,961,605 | 7,961,605 | 7,961,605 | 7,961,605 |
| Final particle images (no.) | 6,732 | 4,314 | 3,185 | 1,682 | 3,400 | 4,204 |
| Map resolution (Å) | 3.79 | 4.12 | 4.22 | 4.57 | 4.16 | 4.16 |
| FSC threshold | 0.143 | 0.143 | 0.143 | 0.143 | 0.143 | 0.143 |

| Construct | Mean (Å) | SD (Å) |
|---|---|---|
| Ligand-free | 18.13 | 1.04 |
| ATP | 17.49 | 0.40 |
| ATP_no_zn2+ | 17.59 | 0.62 |
| ADP | 17.41 | 0.46 |
| AMP | 17.49 | 0.40 |
| cAMP | 17.25 | 0.30 |
| Adenosine | 17.56 | 0.39 |
| Guanosine | 17.8 | 0.75 |
| ATP_R2174A | 17.38 | 0.41 |
| ATP_K2174E | 17.59 | 0.44 |
| ATP_K2584A | 17.46 | 0.42 |
| ATP_K2584E | 17.58 | 0.46 |
| ATP_R2174A_K2177A_K2584A | 17.69 | 0.44 |
| ATP_R2174E_K2177E_K2584E | 17.98 | 0.59 |
| ATP_M2589W | 18.28 | 1.17 |
| ATP_M2589R | 19.36 | 1.00 |

**Supplemental table 2:** Mean and standard deviation for α103-α108 center of mass distance from decorrelated frames.

| Construct | Segment 1 forward primer |
|---|---|
| M2589R | gagcgagcacAACAGATGGCACTACCTG |
| M2589W | GAGCGAGCACAACTGGTGGCACTACCT |
| R2174A, K2177A | CTGACAGCAGAGAGCGCATGTCGCGTCTTC |
| R2174E, K2177E | CTGACAGAAGAGAGCGAATGTCGCGTCTTC |
| K2584A | cgaggagcacATCGCAAGCGAGCACAAC |
| K2584E | cgaggagcacATCGAAAGCGAGCACAAC |

| Segment 1 reverse primer | Segment 2 forward primer |
|---|---|
| tattggtctcCTTAAACCTGTCTTGTAACC | caggtttaagGAGACCAATAGAAACTGGG |
| TCTCTGTCTCGACAAGCCCAGTTTCTATTG | TGGGCTTGTCGAGACAGAGAAGACTCTTGC |
| agcatgcggtACCGGGAGATG | atctcccggtACCGCATGCTAT |
| gcatagcatgcggtACCGGGAGATG | atctcccggtACCGCATGCTATgca |
| acctgtcttgTAACCTTGATACTTACCTGC | atcaaggttaCAAGACAGGTTTAAGGAGACC |
| acctgtcttgTAACCTTGATACTTACCTGC | atcaaggttaCAAGACAGGTTTAAGGAGACC |

| Segment 2 reverse primer |
|---|
| gccaTCTgttGTGCTCGCTCTTGATGTGC |
| GTGCCAGTTGTGCTCGCTCTTGATGTGCTC |
| GCGACATGCGCtctcTGCtgtCAGGTAC |
| GCGACATTCGCtctcTTCtgtCAGGTAC |
| cgctTGCgatGTGCTCCTCGAAGGACAC |
| cgctTTCgatGTGCTCCTCGAAGGACACGG |

**Supplemental table 3:** Oligonucleotides used for site-directed mutagenesis.
